# Functional genomics analysis of developing zebrafish and human endoderm reveals highly conserved *cis*-regulatory modules controlling vertebrate organogenesis

**DOI:** 10.1101/2025.04.08.647835

**Authors:** Daniela M. Figiel, Randa Elsayed, Mark D. Walsh, Simaran Johal, Ying Lin, Till Bretschneider, Sascha Ott, Andrew C. Nelson

## Abstract

While vertebrate species are superficially diverse, they share key commonalities in terms of overall morphology, and organ configuration and function. Maintenance of these traits during evolution is partially explained by conservation of critical genes governing embryonic development. However, for conserved genes to deliver consistent developmental outcomes between species, similar gene regulatory programmes and gene expression patterns must also be maintained. The endoderm germ layer makes major contributions to the respiratory and gastrointestinal tracts, and associated organs including liver and pancreas. We used functional genomics approaches to identify highly conserved endodermal *cis*-regulatory modules (CRMs) functioning across the 400 million years of evolution separating zebrafish and humans. Our analyses suggest that there are few endoderm-specific CRMs, with many CRMs governing pancreas developmental also likely acting within the nervous system. Furthermore, these highly conserved CRMs are strongly enriched for binding sites of “neuro-pancreatic” transcription factors governing both pancreas and nervous system development, potentially suggesting function across these distinct organ systems. Additionally, we identify highly conserved CRMs likely participating endodermal patterning of adjacent craniofacial structures and sensory tissues. The highly conserved CRMs we identify are characterised by conserved patterns of transcription factor binding site co-occurrence. However, they are not characterised by rigid arrangement of binding sites, suggesting more complex or individual grammatical rules. Overall, our analyses provide key insights into critical gene regulatory control during vertebrate endoderm organogenesis, and define a compendium of highly conserved CRMs that should be prioritised for analysis of neuro-pancreatic gene transcriptional control, and anterior embryonic patterning.

## Introduction

While vertebrates exhibit remarkable phenotypic diversity, there are nevertheless many key commonalities of the vertebrate body plan. These include anterior-posterior polarity, similar internal and sensory organs of broadly conserved function, and similar configuration and integration of these organs. Conserved aspects of the vertebrate body plan are accepted to be largely controlled via selective pressures maintaining key gene sequences and functions. Indeed, though the last shared common ancestor of humans and zebrafish is estimated to have inhabited the Earth 400 million years ago, 70% of human genes have at least one obvious zebrafish orthologue (Howe et al. 2013). However, for conserved genes to mediate consistent phenotypic outcomes there must also be consistent regulation of such genes by *cis*-regulatory modules (CRMs). CRMs consist of clusters of transcription factor binding sites (TFBSs) that regulate the expression of target genes through the collective or competitive binding of operative transcription factors (TFs). However, identification of conserved CRMs is often confounded by rearrangement and substitution of TFBSs, leading to similar functional capabilities without deep sequence conservation (reviewed in (Nelson and Wardle 2013; Long et al. 2016; Jindal and Farley 2021). In spite of this, comparative genomics analyses indicate that short discrete non-coding regions of the genome display a high degree of sequence conservation across hundreds of million years of evolution (Elgar 2009; Vavouri and Lehner 2009; Nelson and Wardle 2013; Polychronopoulos et al. 2017). These so-called highly conserved non-coding elements (HCNEs) show non-random distribution in the genome, tending to cluster around genes controlling developmental transitions and cell fate decisions (Sandelin et al. 2004; Woolfe et al. 2005; Engstrom et al. 2008). Many of the products of such genes are transcription factors, hence the putative targets of HCNE CRMs have been referred to as *trans-dev* genes (Woolfe et al. 2005; Elgar 2009; Nelson and Wardle 2013). That such genes appear to be regulated by HCNEs suggests particular constraints on the regulatory logic and architecture on *trans-dev* gene CRMs, though understanding of this is currently lacking.

While the importance and complete range of functions of HCNEs remain to be fully understood, many have been shown to direct tissue-specific gene expression consistent with them acting as enhancers (Nelson and Wardle 2013). Moreover, disruption of specific HCNEs is associated with developmental disorders and diseases including cancer, further highlighting their importance (Polychronopoulos et al. 2017). Functional genomics studies have indicated that sequence conservation at active enhancers, and use of equivalent enhancers between species is maximal during the so-called phylotypic period – a developmental window exhibiting maximal interspecies similarity within a phylum (Von Baer 1828; Duboule 1994; Bogdanovic et al. 2012; Raff 2012; Tena et al. 2014; Martinez-Morales 2016). However, since previous studies typically utilized whole embryos, an analysis of HCNEs within CRMs acting in specific germ layers and tissues across disparate vertebrate species is generally lacking.

Identification of tissue-specific CRMs and their key operative TFs is pivotal to understanding how gene regulatory control of cell fate decisions is achieved during normal development. Furthermore, to understand the basis for developmental disorders, it will be necessary to decipher gene regulatory networks (GRNs) underpinning normal development. Zebrafish is an excellent model organism for dissecting gene regulatory control of cell fate and behaviour (Figiel et al. 2021). While early functional genomics studies analysing whole embryos have been useful, ensemble averaging of signal across all cells leads to both a loss of cell type-specific information, and loss of signal for CRMs functioning in minor cell populations.

The endoderm, one of three primary germ layers of the vertebrate embryo, is induced in early development by Nodal signalling and makes major contributions to the formation of liver, pancreas, intestines, pharynx, swim bladder and other organs (Warga and Nusslein-Volhard 1999; Figiel et al. 2021). The dorsal forerunner cells (DFCs) which go on to form the zebrafish organ of laterality, Kupffer’s vesicle (KV) are also induced by Nodal and are suggested by some to be a specialized dorsal subset of endodermal cells due to their similar early developmental program (Alexander and Stainier 1999; Warga and Kane 2018; Moreno-Ayala et al. 2021). The SOX family transcription factor Sox17 is a commonly used marker of endoderm across vertebrate species, its expression indicating definitive specification of endoderm during gastrulation (Hudson et al. 1997; Alexander and Stainier 1999; Kanai-Azuma et al. 2002). As well as early expression in endoderm progenitors, *sox17* is subsequently expressed in other progenitor populations and has key roles in blood formation, vasculature development and also left-right patterning throughout vertebrate evolution (e.g. (Kanai-Azuma et al. 2002; Aamar and Dawid 2010; Chung et al. 2011; Saund et al. 2012; Viotti et al. 2012; Lilly et al. 2017; Figiel et al. 2021; Johal et al. 2024). While multiple recent studies have sought to identify CRMs and gene expression patterns in lineages derived from *sox17*-expressing progenitors (Quillien et al. 2017; Bonkhofer et al. 2019; Dobrzycki et al. 2020; Lopez-Perez et al. 2021; Xia et al. 2021; Trinh et al. 2023), substantial gaps persist in our knowledge of gene regulation in the developing endoderm. *Tg(sox17:GFP)* reporter zebrafish (hereafter referred to as *sox17:GFP*) express GFP in all endodermal cells throughout early organogenesis stages, so can be used to isolate endodermal cells (Sakaguchi et al. 2006; Mizoguchi et al. 2008; Fukui et al. 2014). However, expression of *sox17* in other lineages limits utility of *sox17:GFP* in isolating a pure endoderm population without the use of other reporter transgenes. We have combined *sox17:GFP* with red fluorescent reporters of endothelial and erythroid lineages to enrich for endoderm cells during early and mid-organogenesis stages –

28 and 48 hours post-fertilisation (hpf). We have followed this with ATAC-seq to identify accessible chromatin regions potentially acting as CRMs. We have integrated these data with published datasets from human embryonic stem cells (hESCs) that have undergone directed differentiation to represent distinct endoderm cell populations along the anterior-posterior axis. Our functional genomics analyses reveal over a thousand discrete HCNEs bearing the hallmarks of CRM functionality in both human and zebrafish endoderm. Furthermore, genes proximal to these HCNEs are highly enriched for developmental phenotypes in both animal models and human patients. However, while analyses of the HCNEs themselves reveals strong enrichment for TFBSs consistent with known regulators for endodermal organ formation, we find only limited evidence for consistent arrangement of TFBSs across endodermal HCNEs within species.

Overall our analyses reveal a compendium of HCNE CRMs functioning in endodermal tissues likely to be governing consistent aspects of the vertebrate body plan, which should be prioritised for further functional dissection and investigation at scale.

## Results

### ATAC-seq reveals CRMs functioning in distinct sox17-expressing lineages during zebrafish embryogenesis

To enrich for endodermal cells during organogenesis we exploited *sox17:GFP* fish (Mizoguchi et al. 2008). While endogenous *sox17* is rapidly downregulated at the end of gastrulation (Alexander and Stainier 1999), GFP protein persists throughout the endoderm for days after the endogenous gene has been silenced (Fig. 1A). However, *sox17* expression is also observed in erythroid and endothelial lineages, limiting the utility of *sox17:GFP* alone for enrichment of endoderm (Chung et al. 2011). We therefore crossed homozygous *sox17:GFP* fish with fish homozygous for both *gata1a:dsRed* and *kdrl:mCherry* transgenes (Fig. 1B). We then used fluorescence activated cell sorting (FACS) to separate *sox17+* vascular and erythroid mesodermal lineages (termed *sox17*M) from endoderm-enriched *sox17*+ lineages (*sox17*E) and all other *sox17-* lineages (*sox17*N) at 28 and 48 hpf (Fig. 1C). 28 hpf was chosen as the earliest timepoint to ensure robust detection of all three fluorescent proteins. To verify that our sorting strategy enriched for endodermal cells in the *sox17*E population we performed bulk RNA-seq on 28 hpf samples. This revealed strong enrichment for transcripts expressed in specific endoderm cell populations including pancreatic, liver and intestinal markers (Fig. 1D, Supplemental Table S1). However, we also note enrichment of markers of fin epithelia and notochord which we attribute to transdifferentiation of the *sox17*-expressing left-right organizer (Kupffer’s vesicle - KV), which gives rise to posterior cell types once its role is complete ((Ikeda et al. 2022), Supplemental Fig. S1). Nevertheless, the *sox17*E population shows strong enrichment for endoderm markers, thus validating our approach. We therefore proceeded to perform independent duplicate ATAC-seq analysis on the three distinct FACS populations at 28 and 48 hpf (Supplemental Fig. S2).

**Figure 1.**
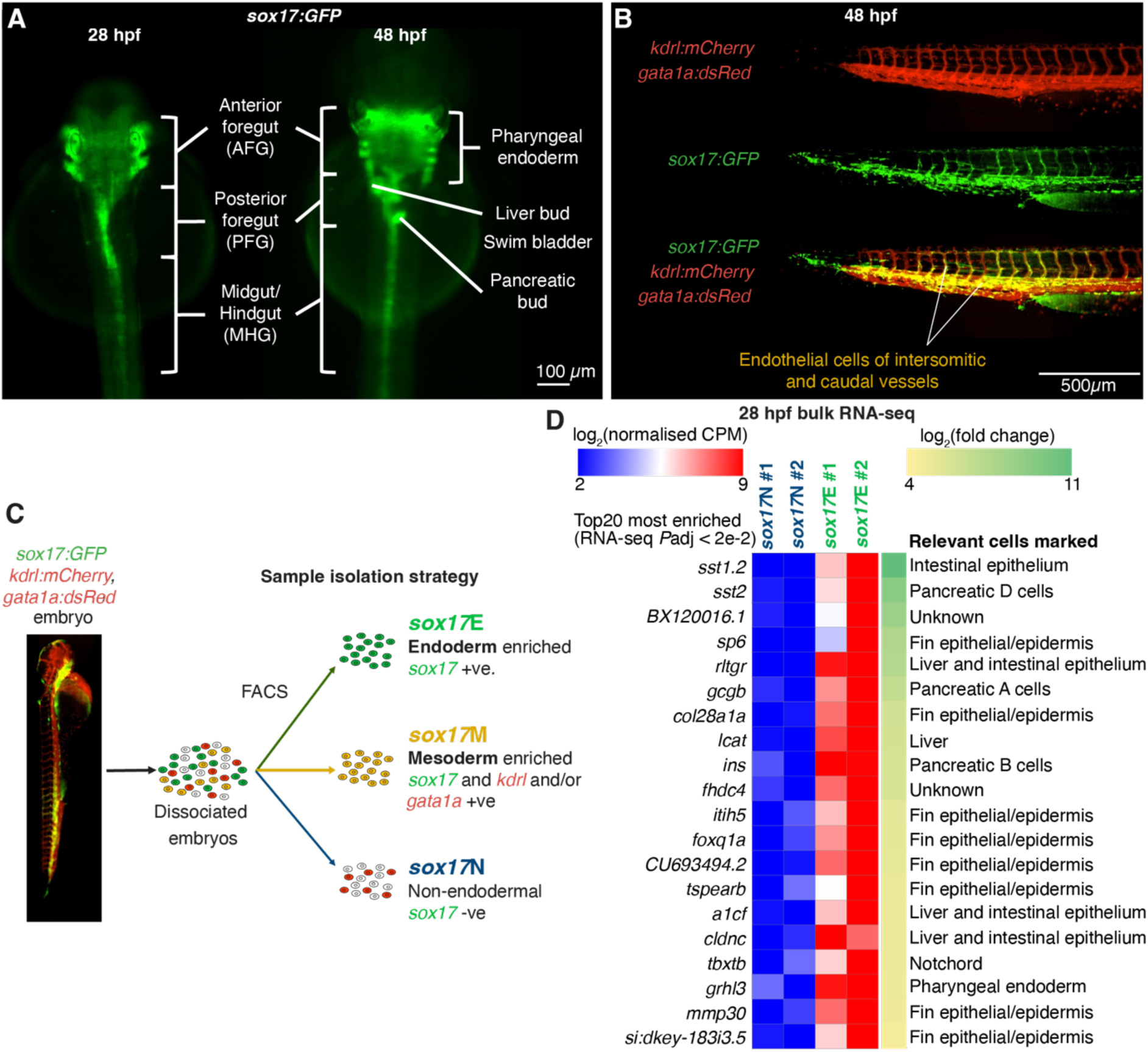
Fluorescence activated cell sorting separates endodermal and mesodermal cell populations arising from *sox17+* progenitors. (A) Widefield images of *sox17:GFP* embryos at 28 and 48 hpf with distinct endodermal structures indicated. (B) Widefield images of coexpression of *sox17:GFP* with *kdrl:mCherry* and *gata1a:dsRed* transgenes in trunk and tail vasculature at 48 hpf. (C) Sorting strategy. (D) RNA-seq heatmap of the top 20 significant genes showing strongest fold enrichment in *sox17*E over *sox17*N at 28 hpf.

ATAC-seq analysis revealed thousands of differentially accessible regions (DARs) distinguishing the three sorted populations both within and between timepoints (Fig. 2A,B, Supplemental Tables S2,3). Chromatin accessibility profiles were consistent with the predicted cell identities being enriched in each sorted cell population. For example, the embryonic haemoglobin gene cluster, erythroid marker *gata1a*, and endothelial marker *fli1b* show enhanced accessibility in the *sox17*M population (Fig. 2C), while markers of the posterior foregut (*gata6* and *foxa3*), pancreas and duodenum (*pdx1*), and liver and intestine (*fabp2*) show enhanced accessibility in the *sox17*E population (Fig. 2D). However, we note that many more regions show enhanced accessibility in *sox17*E over *sox17*M, compared to *sox17*E over *sox17*N (Fig. 2A). This suggests that while there are differences in accessibility profiles that distinguish the *sox17*E population from *sox17*M, many regions of accessible chromatin in *sox17*E are nevertheless shared with cell types in the *sox17*N population rather than being unique to endoderm.

**Figure 2.**
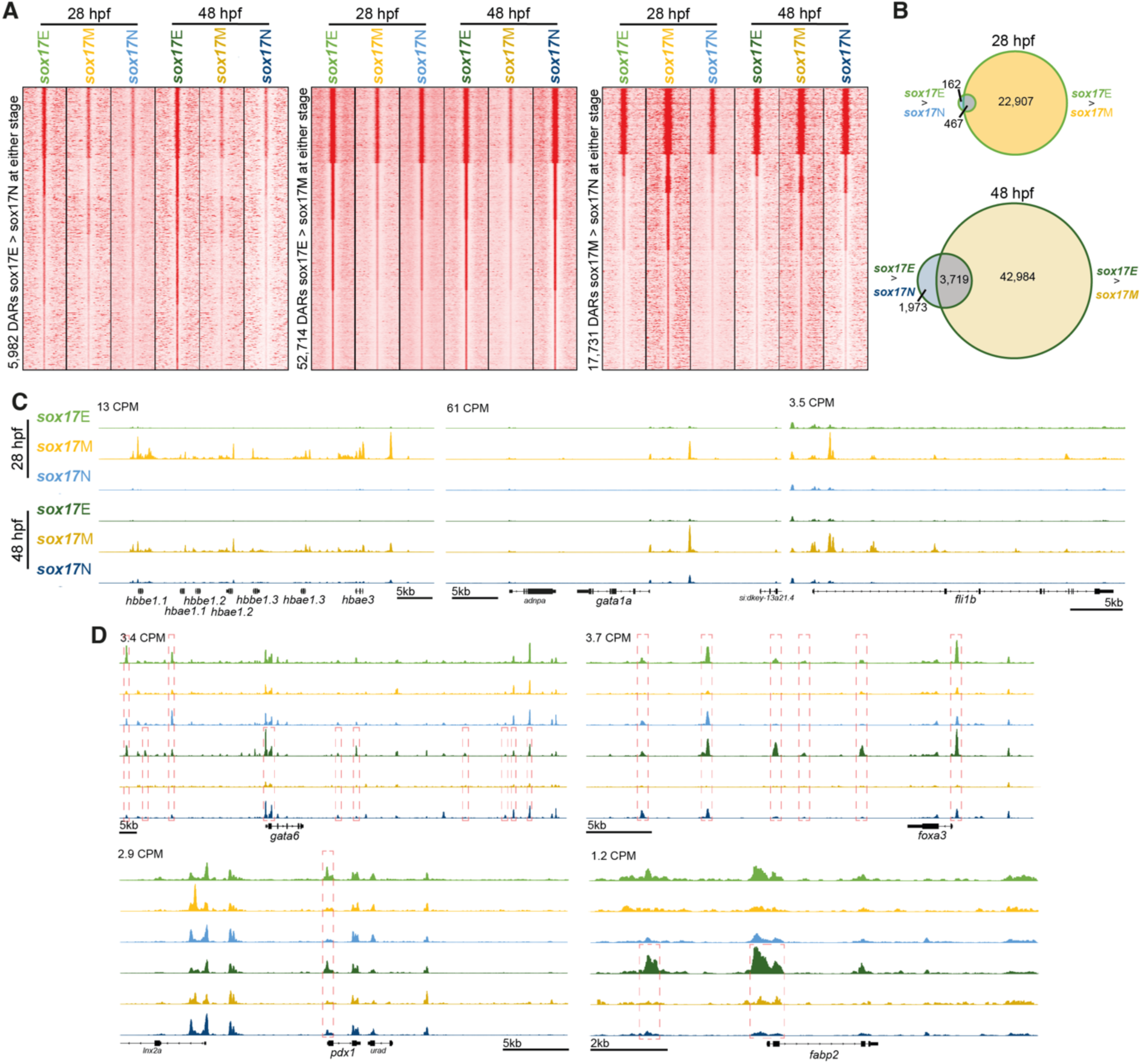
Sorted cell populations have distinct chromatin accessibility profiles indicative of constituent cell identities. (A) Heatmaps of relative ATAC-seq read densities at differentially accessible regions (DARs, FDR ≤ 0.05) for the comparisons indicated. (B) Venn diagrams indicating number of *sox17*E > *sox17*N DARs overlapping *sox17*E > *sox17*M DARs at 28 and 48 hpf. (C) Example loci indicating greater accessibility of erythrocyte marker genes (*gata1* and haemoglobin gene cluster) and the endothelial marker *fli1b* in the *sox17*M population. (D) Example loci indicating greater accessibility of markers of the posterior foregut (*gata6* and *foxa3*), pancreas and duodenum (*pdx1*), and liver and intestine (*fabp2*) in the *sox17*E population compared to *sox17*M and *sox17*N. Peak heights in counts per million reads (CPM) are indicated. Significant DAR comparisons in panel D and outlined in red.

To more globally ascertain whether the ATAC-seq profiles of the sorted populations are consistent with the expected constituent cell populations we performed anatomical enrichment analysis on promoter regions that showed enhanced accessibility in the *sox17*E population. We specifically focused on promoter regions to avoid errant annotation of DARs to putative target genes. At the stages studied the endoderm predominantly consists of epithelial cells of the pharyngeal endoderm and intestinal rod, as well as the developing pancreas and liver primordia. As expected, we found strong enrichment for anatomical terms consistent with epithelial structures, gut, liver and pancreas, for DARs more accessible in both *sox17*E>*sox17*M and *sox17*E>*sox17*N, especially at 48 hpf (Fig. 3A, Supplemental Tables S4-7). This is consistent with greater development of endodermal organs by this stage. We also found enrichment for notochord markers in *sox17*E>*sox17*M, consistent with transdifferentiated cells from KV incorporating into this structure (Supplemental Figs S1,4). Conversely, we found strong enrichment for anatomical terms consistent with nervous system and eye development but not endodermal tissues for *sox17*N>*sox17*E promoter DARs (Supplemental Table S8). This is highly consistent with these structures being absent from the *sox17*E population, as expected. However, remarkably we also found strong enrichment for markers of neural structures in *sox17*E>*sox17*M, and the lateral line system in both *sox17*E>*sox17*M and *sox17*E>*sox17*N (Fig. 3A). This suggests that either cells arising from *sox17*+ endodermal progenitors and KV in the *sox17*E population exhibit similar gene accessibility (and consequent potential expression) signatures to the nervous and lateral line systems, or there is hitherto unrecognized *sox17:GFP* expression in these cell types.

**Figure 3.**
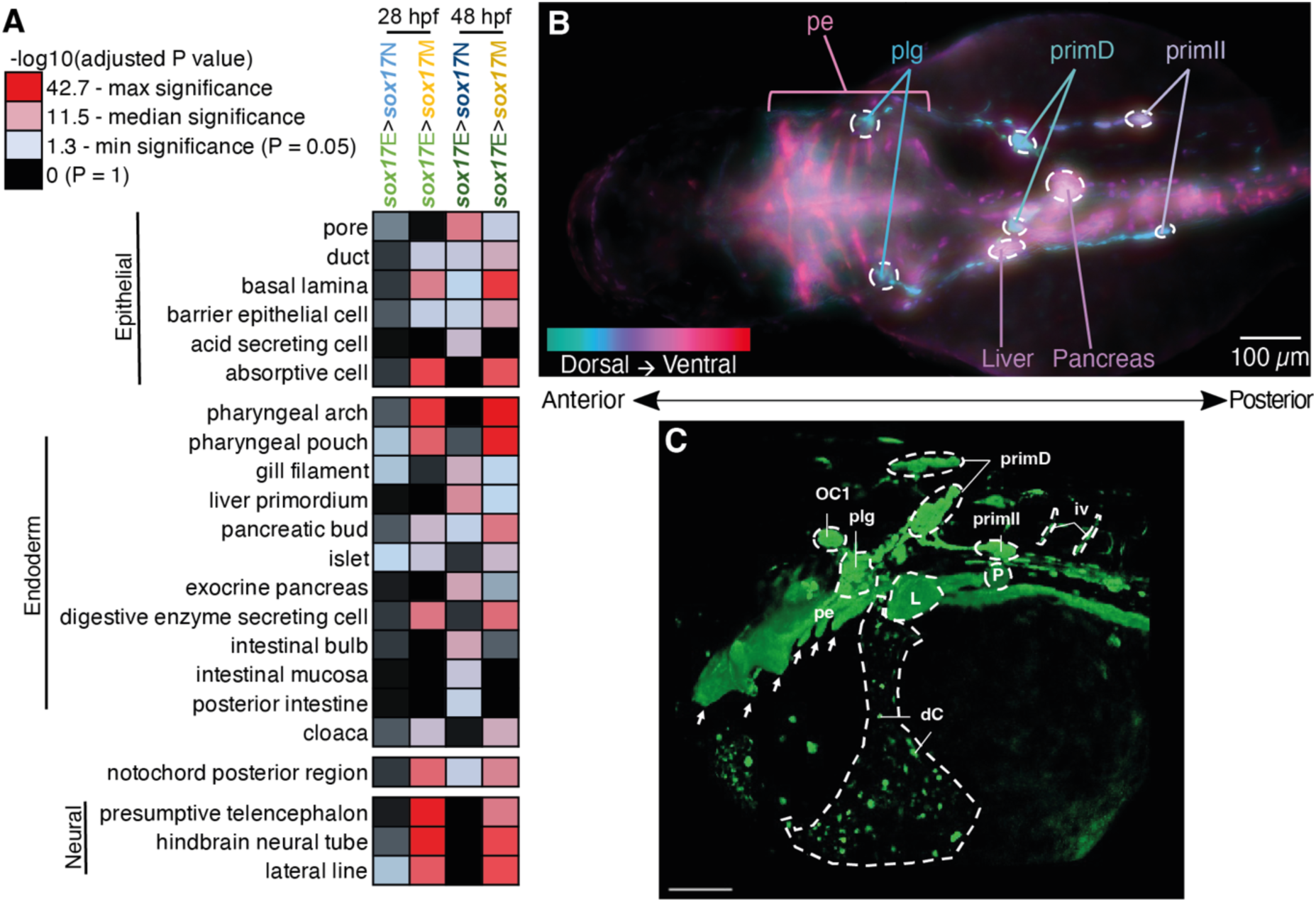
Promoter accessibility within the *sox17*E populations is consistent with presence of endodermal cell populations and lateral line neurons. (A) Heatmap of-log_10_(adjusted P value) from fishEnrichr Anatomy GeneRIF Predicted Z-score analyses of promoters showing greater accessibility in the comparisons indicated (FDR ≤ 0.05) as annotated by ChIPseeker. Selected terms are shown; complete fishEnrichr outputs are in Supplemental Tables S9-10. (B) Dorsal widefield images of *sox17:GFP* embryo at 48 hpf. Nine focal layers have each been artificially coloured based on depth from dorsal to ventral as indicated. (C) 3D rendering of confocal images 48 hpf *sox17:GFP* laterally orientated embryo with head to the left and tail to the right. Scale bar = 100 µm. OC1 = organ of Corti 1, plg = posterior lateral line ganglion, primD = dorsal primordium, primII = second primordium, iv = intersegmental vessel, L = liver, P = pancreas, dC = cells within duct of Cuvier, pe = pharyngeal endoderm. Pharyngeal pouches are marked by white arrows.

To further explore whether there is GFP expression in the nervous and lateral line systems we performed detailed imaging of *sox17:GFP* embryos. This revealed subtle but detectable expression in a small set of neurons exhibiting morphology consistent with being a subset of lateral line neurons within the hindbrain and trunk (Fig. 3B,C). This includes organ of Corti 1 (OC1), posterior lateral line ganglion (plg), dorsal primordium (primD) and second primordium (primII) (Pujol-Marti and Lopez-Schier 2013). We otherwise did not detect GFP expression in developing neural structures. Development of the posterior lateral line ganglion, nerve and its projections have been shown to be affected by inhibition of retinoic acid (RA) signalling (Begemann et al. 2004). Consistent with expectations, we found that modulation of RA signalling through addition of RA itself, an inhibitor of *Aldh1a* mediated RA synthesis (diethylaminobenzaldehyde – DEAB), and an inverse agonist of RA Receptor (BMS493) all disrupted the formation of these *sox17:GFP*+ neurons (Supplemental Fig. S5). We therefore conclude that the *sox17:GFP* reporter is expressed in a subset of lateral line neurons, but not other neural structures.

Overall we conclude that we have profiled chromatin accessibility across two distinct *sox17*-expressing populations – *sox17*M containing erythroid progenitors and a subset of endothelial cells, and *sox17*E containing all endoderm, plus KV-derived posterior cell types and lateral line neurons. Given *sox17*E captures endodermal chromatin accessibility and thus active CRMs, we next wanted to determine which of these CRMs are likely to act during endoderm formation across the 400 million years of evolution separating zebrafish and humans.

### Highly conserved CRMs functioning in both human and zebrafish endoderm cluster around genes controlling diverse aspects of endodermal and vertebrate-specific development

To determine which potential CRMs captured by our ATAC-seq data are recognisably functional in human endoderm we compared our zebrafish data to existing ChIP-seq data for the marker of active promoters and enhancers, H3K27ac (Creyghton et al. 2010). Specifically, we utilised H3K27ac ChIP-seq data from human embryonic stem cells (hESCs) that had undergone efficient directed differentiation to either anterior foregut (AFG), posterior foregut (PFG) or midgut/hindgut (MHG) (Loh et al. 2014) (Supplemental Table S11). AFG principally gives rise to anterior structures including thyroid and lungs in human, PFG to liver and pancreas, and MHG to small and large intestine.

We wanted to avoid eliminating DARs that are potentially functionally important in the endoderm, while enriching for genomic regions that are not likely to be controlling constitutively expressed genes. We therefore primarily focused our attention on DARs showing greater accessibility in *sox17*E relative to *sox17*M (*sox17*E>*sox17*M DARs). This is because our initial analyses indicate that relatively few DARs distinguish *sox17*E from the *sox17*N population, but that the majority that do also distinguish *sox17*E from *sox17*M (Fig. 2A,B).

To identify zebrafish *sox17*E DARs corresponding to human H3K27ac ChIP-seq peaks we compared both datasets to highly conserved non-coding elements (HCNEs) exhibiting ≥70% sequence identity across ≥30 alignment columns (Engstrom et al. 2008). We identified a total of 5,000 HCNEs overlapping H3K27ac peaks in at least one of the AFG, PFG and MHG cell populations (Fig. 4A). 7,435 HCNEs overlap *sox17*E>*sox17*M DARs while 236 HCNEs overlap *sox17*E>*sox17*N DARs, of which 112 are common to both *sox17*E>*sox17*M and *sox17*E>*sox17*N DARs (Fig. 4B). Of the 5,000 HCNEs overlapping human H3K27ac peaks, 1,701 were also identified in *sox17*E>*sox17*M DARs (Fig. 4C,D, Supplemental Fig. S6-7, Table S12-13). These HCNEs occur proximal to genes with prominent roles in endoderm development such as regulator of gut development *DLL1*/*dld* (Pellegrinet et al. 2011; Troll et al. 2018) (Fig. 4E), regulator of pancreas development *FOXA2*/*foxa2* (Shin et al. 2008; Lee et al. 2019; Elsayed et al. 2021) (Fig. 4F) and pancreatic and biliary regulator *HES1*/*her6* (Fig. 4G) (Spence et al. 2009).

**Figure 4.**
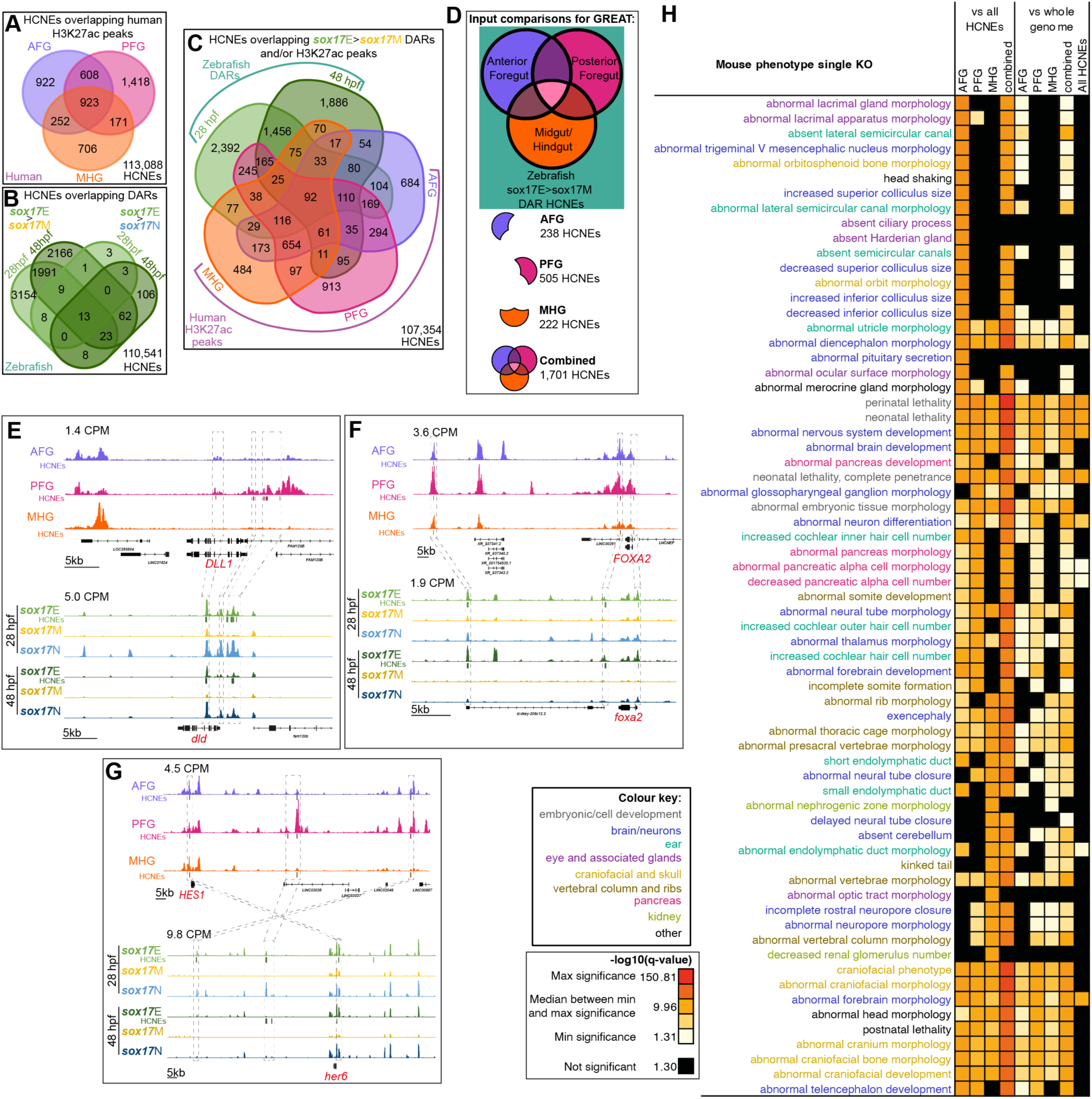
HCNE CRMs bearing the hallmarks of functionality in human and zebrafish endoderm are proximal to genes with deleterious pancreatic, neural and craniofacial phenotypes. (A) Number of HCNEs overlapping human H3K27ac peaks in AFG, PFG and/or MHG patterned endoderm cell populations derived from hESCs. Number of HCNEs not overlapping H3K27ac peaks are at the bottom right. (B) Distribution of HCNEs overlapping *sox17*E>*sox17*M DARs vs *sox17*E>*sox17*N DARs at 28 and 48 hpf. Number of HCNEs not overlapping DARs are at the bottom right. (C) Distribution of HCNEs overlapping *sox17*E>*sox17*M DARs also overlapping human H3K27ac peaks in AFG, PFG and/or MHG patterned endoderm cell populations derived from hESCs. Number of HCNEs not overlapping H3K27ac peaks or DARs are at the bottom right. (D) Schematic of human HCNE subsets used for functional, anatomical and phenotypic enrichment analyses using GREAT. (E-F) Example orthologous loci bearing zebrafish HCNE DARs and human H3K27ac HCNE peaks. Normalised peak heights in counts per million reads are indicated. HCNEs overlapping peaks and DARs are displayed beneath and colour-coded as the tracks. Grey boxes and lines indicate regions with HCNE matches between species. (H) Heatmap of enrichment of single gene mouse knockout phenotypes for collections of HCNEs as indicated in panel D. Significance in-log10(HyperFdrQ) are indicated both relative the whole genome, and also to all HCNEs to account for non-random distribution of HCNEs. Terms are colour-coded according to anatomical categories as indicated.

To more globally analyse the targets of putative CRMs acting in both human and zebrafish endoderm we used GREAT to perform functional and phenotypic annotation analyses on the common 1,701 HCNEs (McLean et al. 2010). To determine whether the HCNEs acting in AFG, PFG and MHG endoderm show distinct characteristic distribution patterns indicative of alternative target genes we analysed subsets unique to each population as well as the complete combined set (Fig. 4D). Furthermore, since HCNEs are known to have non-random genomic distribution patterns with specific biases towards *trans-dev* genes we tested whether the 1,701 HCNEs (hereafter referred to as “endoderm HCNEs”) showed enrichment around specific classes of gene both overall and compared to all HCNEs (Fig. 4H, Supplemental Fig. S8-12).

Consistent with expectations, our analyses reveal a highly significant association between endoderm HCNEs and genes encoding DNA binding and gene regulatory proteins, including HMG domain-containing and chromatin binding factors. Notably, however, the endoderm HCNEs appear to show a greater enrichment of such genes compared to all HCNEs (Supplemental Fig. S8). Furthermore, both Biological Process Gene Ontology terms, and mammalian phenotype terms associated with endoderm HCNE subsets from AFG, PFG and MHG cells are broadly consistent with their anterior-posterior location within the vertebrate body plan. For example, putative target genes of HCNEs shared between AFG H3K27ac peaks and *sox17*E DARs are significantly associated with GO Biological Process Terms, mouse phenotypes and human phenotypes focused on anterior structures including brain, ear and craniofacial structures (Fig. 4H, Supplemental Figs S9,11,12). Similarly, putative targets of PFG H3K27ac peaks and *sox17*E DARs are associated with pancreas development and pancreatic abnormalities in knock-out mice, and also notably cardiac defects in humans (Fig. 4H, Supplemental Fig. S12). Conversely, putative targets of MHG H3K27ac peaks and *sox17*E DARs appear to be largely associated with formation of tube and ductal structures such as neural tube closure, endolymphatic duct and kidney formation, as well as vertebral column formation (Fig. 4H, Supplemental Figs S9,11,12). This may be indicative of common gene regulatory programmes governing tube formation including the gut, and CRM accessibility potentially being broadly coordinated regionally along the anterior-posterior axis.

We note that many of the putative targets of HCNE CRMs identified in human and zebrafish endoderm function in nervous system development (Fig. 4H, Supplemental Figs S9,11,12). For example, *FOXA2*/*foxa2* functions both in pancreas development and formation of the floor plate of the neural tube (Ang and Rossant 1994; Brand et al. 1996; Shin et al. 2008; Dal-Pra et al. 2011). Similarly, *HES1*/*her6* has been shown to exhibit nervous system defects including premature neurogenesis, severe neural tube defects, increased numbers of pulmonary neuroendocrine cells, and also pancreatic hypoplasia in mouse knock-out models (Ishibashi et al. 1995; Ito et al. 2000; Jensen et al. 2000), and regulates cell proliferation in the hindbrain in zebrafish (Coolen et al. 2012). As we discuss later, it is therefore highly likely that the significant enrichment for terms associated with the nervous system can be attributed to common neuro-pancreatic gene regulatory programmes operating across the developing nervous system and pancreas. Overall we conclude that we have identified a compendium of HCNEs likely to control gene expression during early organogenesis across vertebrate evolution.

### HCNEs bearing hallmarks of functionality in human and zebrafish endoderm are enriched for binding sites of endoderm transcription factors

To determine which transcription factors are likely to be acting via the endodermal HCNEs we performed *de novo* and known TFBS enrichment analysis in the human and zebrafish sequences (Fig. 5A-C). We particularly focused on the PFG population given its role in pancreas development and the strong association between putative HCNE target genes and pancreas development in our previous analyses (Fig. 4H). Since TFBSs are often degenerate, with multiple TFs potentially able to bind the same site, we also analysed expression of the candidate TFs. This revealed notable concordance between enrichment of some TFBSs and expression of the TF (Fig. 5C). For example, TFBSs for SOX21 and MEIS1 show greater enrichment in AFG and PFG HCNEs than MHG HCNEs, and *SOX21* and *MEIS1* are also more highly expressed in AFG and PFG. Similarly, HOXA11, HOXD11 and HOXA13 TFBSs show greater enrichment in MHG HCNEs, consistent with greater expression of the TFs in MHG endoderm. However, we also find strong enrichment for TFBSs of multiple TFs known to have key functions in the endoderm in spite of the TFs showing no/minimal expression in AFG, PFG or MHG RNA-seq datasets (Fig. 5C). For example, ASCL1, ASCL2, PTF1A, PDX1, NEUROD1 and NKX6.1 are all known or suggested to have roles in pancreatic endoderm development (Offield et al. 1996; Naya et al. 1997; Krapp et al. 1998; Sander et al. 2000; Yee et al. 2001; Kawaguchi et al. 2002; Dong et al. 2008; Binot et al. 2010; Flasse et al. 2013; Duque et al. 2022; Vanheer et al. 2023; Zhu et al. 2023) and their TFBSs are enriched in PFG HCNEs though their expression is not appreciably detected in PFG endoderm. However, we find strong expression of these TF genes in RNA-seq data from hESCs further differentiated beyond the PFG stage to pancreatic progenitors (PPs). This suggests that the HCNE CRMs these TFs act through bear functional marks prior to the expression of these TFs. Notably, multiple SOX family transcription factors including SOX2, 3, 4, 6, 9, 15, 17 and 21 all show enrichment in endoderm HCNEs and appreciable expression in PFG and PP populations, as do FOXA1 and FOXA2. Given prior evidence that SOX and FOX transcription factors often have pioneer activity (Kamachi and Kondoh 2013; Iwafuchi-Doi et al. 2016; Julian et al. 2017; Fuglerud et al. 2022), it is tempting to speculate that earlier expression of these TFs may render endodermal HCNEs accessible for subsequent binding by later expressed pancreatic TFs like PTF1A.

**Figure 5.**
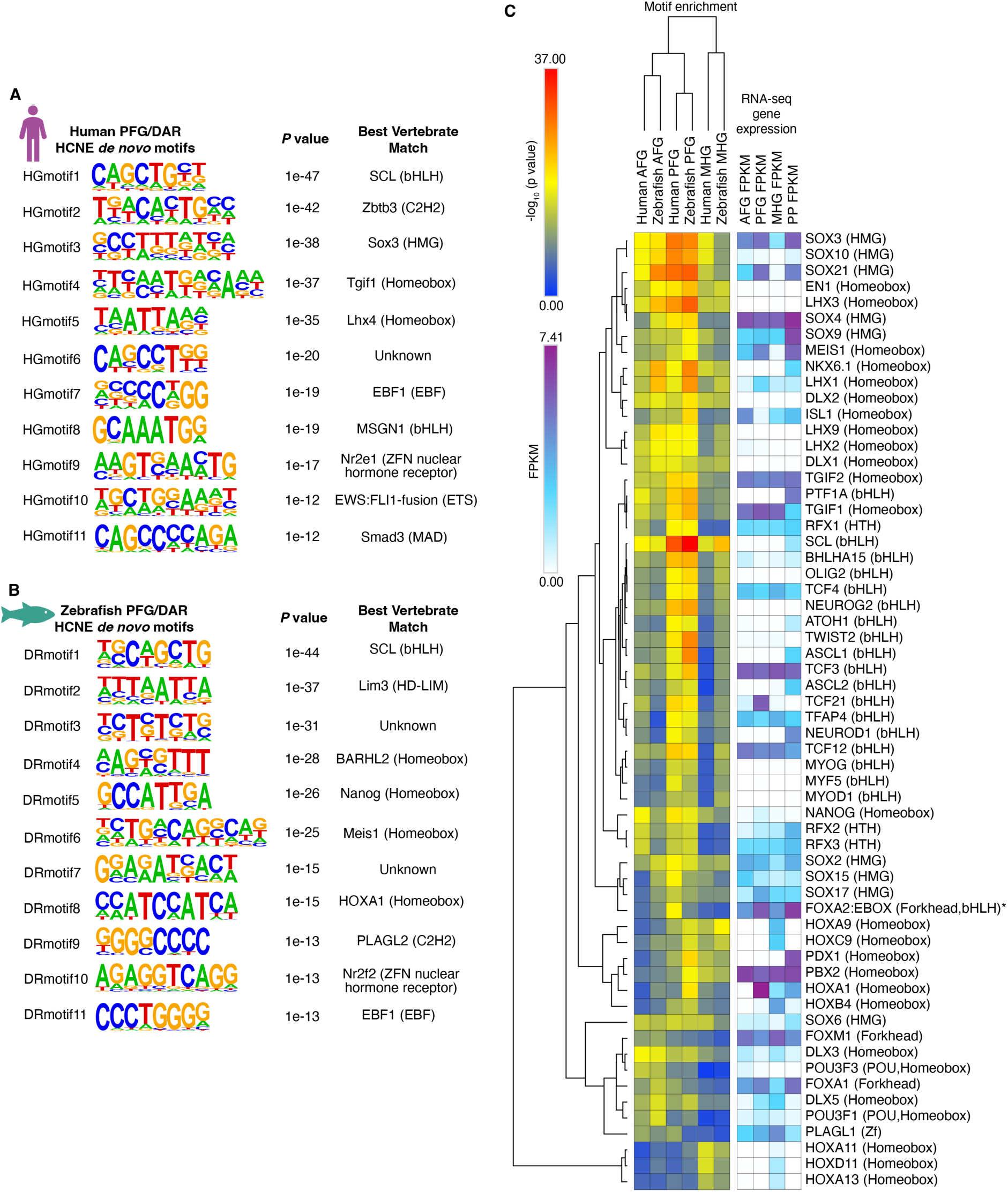
Motif enrichment analysis reveals candidate transcription factors acting via endoderm HCNE CRMs. (A-B) High confidence *de novo* motifs enriched in HCNEs overlapping both human PFG H3K27ac peaks and *sox17*E>*sox17*M DARs at 28 and/or 48 hpf, for the human (A) and zebrafish (B) HCNE sequences. *P* vales and closest match vertebrate transcription factors assigned by HOMER are indicated. (C) Hierarchical clustering of enrichment of known motifs in human and zebrafish endoderm HCNEs overlapping both *sox17*E>*sox17*M DARs at 28 and/or 48 hpf, and human AFG, PFG and MHG H3K27ac peaks as indicated. Enrichment in both human and zebrafish HCNE sequences are shown. RNA-seq expression of transcription factor genes corresponding to enriched motifs is indicated as FPKM (Fragments Per Kilobase of transcript per Million mapped reads). AFG = anterior foregut; PFG = posterior foregut; MHG = mid/hindgut; PP = pancreatic progenitors. * While the enriched motif is FOXA2:EBOX FPKM heatmap indicates *FOXA2* expression alone.

### PFG HCNEs have characteristic patterns of TFBS co-occurrence, but show limited evidence for consistent rigid grammatical constraint

Given we identify TFBSs of major regulators of pancreas development in PFG HCNEs identified in both species, we conclude that these highly conserved putative CRMs are likely to be important for pancreas development. That the HCNE sequences are conserved between humans and zebrafish demonstrates that arrangements of putative TFBSs have remained largely consistent across 400 million years of evolution. Constrained TFBS “grammar” (arrangement, spacing and orientation of TFBS) has historically been suggested to point to the “enhanceosome” model of CRMs, where rigid TFBS grammar is required for correct assembly of TF complexes (reviewed in (Long et al. 2016; Jindal and Farley 2021). We wanted to determine whether PFG HCNEs contain consistent sets of grammatically constrained TFBSs, potentially suggestive of consistent TF complexes acting across subsets of HCNEs. We therefore analysed both co-occurrence of the identified enriched TFBSs within HCNEs, and also whether co-occurring TFBSs showed significantly consistent spacing patterns (Fig. 6).

**Figure 6.**
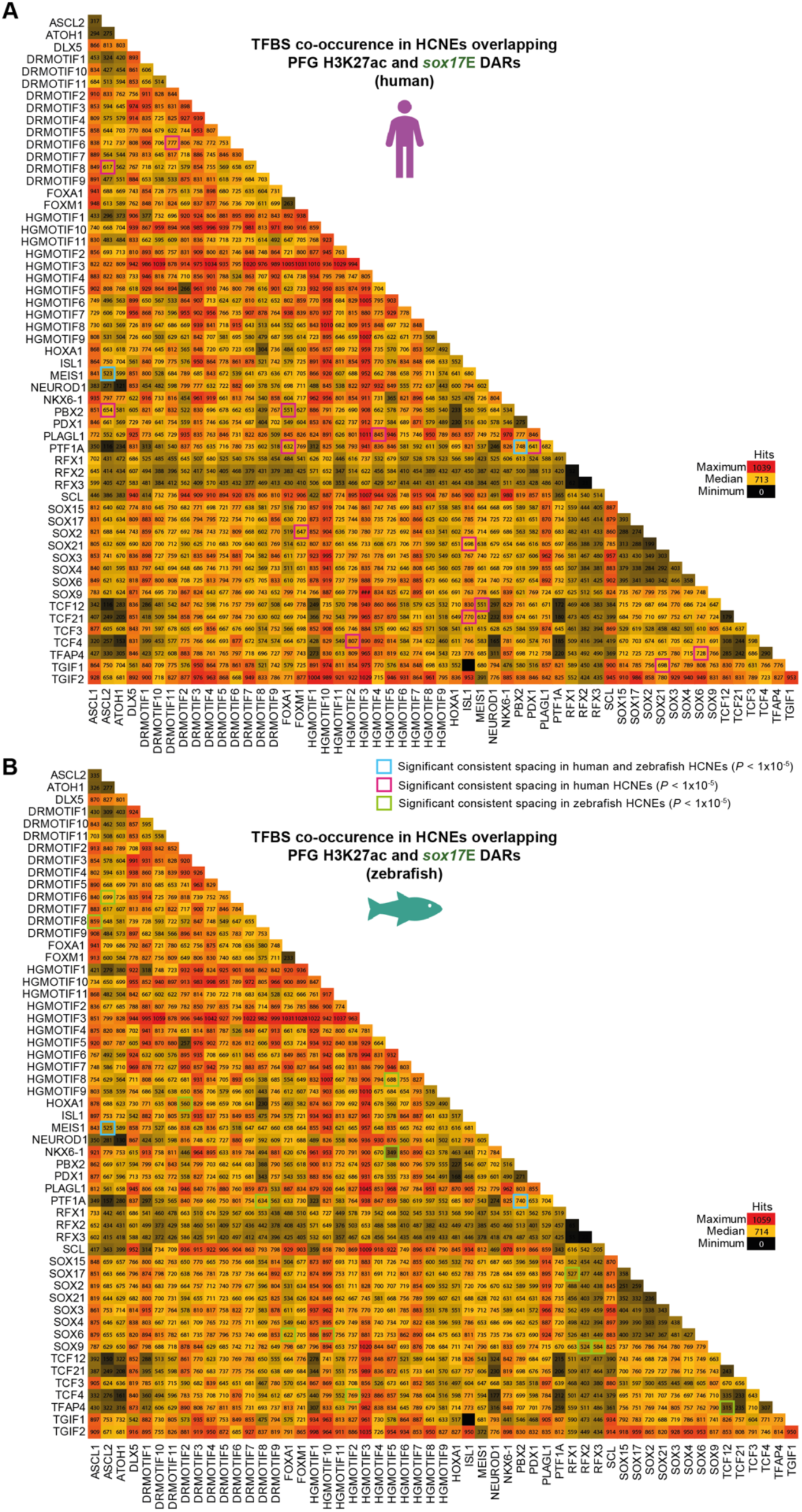
Human and zebrafish endoderm HCNEs show consistent coincidence of transcription factor binding sites (TFBSs) h limited grammatical consistency either within or between species. (A) Heatmap matrix indicating the number of human s overlapping both human PFG H3K27ac peaks and *sox17*E>*sox17*M DARs at 28 and/or 48 hpf containing a significant match *P* value < 0.05) for motif pairs as indicated. (B) As panel A but for zebrafish HCNE sequences. Co-occurring motifs showing antly consistent spacing (SpaMo *P* value < 1 x 10^-5^) are outlined and colour-coded to represent whether significantly consistent g is found in human or zebrafish HCNEs or both.

There are well-characterised examples of “suboptimisation” of TFBSs within developmental CRMs, where low affinity TFBSs tune target gene expression preventing deleterious ectopic or overexpression (Farley et al. 2015; Farley et al. 2016). However, position frequency matrices used for most motif analyses are derived from binding experiments (e.g. ChIP-seq) that are most indicative of high affinity TFBSs. To avoid exclusion of potentially important low affinity TFBSs in our analyses, we applied a permissive cut-off for identification of individual TFBSs. Our results indicate strong pairwise co-occurrence of specific TFs throughout PFG HCNEs, while some TFBS combinations are rare (Fig. 6). As expected, patterns of TFBS co-occurrence are virtually identical between human and zebrafish PFG HCNEs. This suggests that the same sets of TFs are likely capable of acting via the same HCNE CRMs in both species. However, this merely indicates whether TFBSs consistently occur in the same HCNEs, but not whether they show consistent spacing or orientation. We therefore next tested whether TFBS pairs have consistent arrangement and spacing, potentially indicative of constrained binding of TF complexes. While we find 16 TF pairs showing consistent arrangement in human PFG HCNEs and 15 in the homologous zebrafish HCNEs (*P* < 1 x 10^-5^), only PTF1A-PBX2 and MEIS1-ASCL2 were identified in both species (Fig. 6A,B). Notably, the significant spacing of PTF1A-PBX2 and MEIS1-ASCL2 TFBSs was the same in both species (12 and 12 bp respectively).

Overall our analyses reveal consistent patterns of TFBS co-occurrence in PFG HCNEs between humans and zebrafish, potentially indicative of conserved cooperative gene regulatory programmes. However, there is limited evidence for highly consistent CRM grammar both across sets of HCNEs and between species. This potentially suggests that either few of these TFs act in complexes with each other, or the complexes are flexible in terms of combined recognition of target TFBSs.

### HNF1B HCNE CRMs identified in endoderm drive expression in the developing hindbrain

We next wanted to determine whether the HCNE CRMs we identified in human and zebrafish endoderm are capable of driving expression in the predicted endodermal tissues. We selected *HNF1B* and its zebrafish orthologue *hnf1ba* as a target gene of interest due to its key conserved role in pancreas development, and monogenic diabetes in human patients (Fajans et al. 2001; Sun and Hopkins 2001; Quilichini et al. 2021). Our *sox17*E ATAC-seq data indicate regions of accessibility containing HCNEs in both introns 4 and 5, while the homologous human HCNEs are within PFG H3K27ac peaks in the same introns. There is also a *sox17*E ATAC-seq peak 3 kb upstream of the transcription start site that does not harbour HCNEs. To determine whether these putative enhancer regions, and the HCNEs are capable of driving expression in foregut endoderm we tested their ability to drive mCherry reporter expression in *sox17:GFP* embryos. We tested the ability of multiple genomic regions to drive expression. For intron 4 this included the entire accessible zebrafish region (i4Enh +6-8kb), discrete elements on the flanks of the intron 4 accessible region lacking HCNEs (i4Enh +6kb and i4Enh +8kb), and just the HCNE cluster (i4zHCNE). We also similarly tested the equivalent human intron 4 HCNE cluster (i4hHCNE). For intron 5 we tested discrete regions containing the zebrafish and human HCNE (i5zHCNE and i5hHCNE respectively). We also tested the upstream accessible region (Enh-3kb). Each reporter construct also contained the *crystallin, alpha a* (*cryaa*) promoter upstream of *gfp*. This drives GFP expression in the lens of the eye, thus providing a constitutive marker to control for injection. We imaged the zebrafish regularly up to and including 48 hpf, consistent with the terminal ATAC-seq timepoint. Except for Enh-3kb, all constructs drove expression in multiple tissues, with the most consistent mCherry expression seen in hindbrain (Fig. 7B,C, Supplemental Figs S13-14, Tables S14-15). None of the reporter constructs yielded mCherry co-expression with *sox17:GFP*, indicating that they are not individually capable of driving expression in the endoderm (or lateral line neurons). All constructs containing HCNEs drove expression in the developing hindbrain (i4zHCNE – 62.2% of embryos; i4hHCNE – 87.5%; i4Enh +6-6kb 28.6%; i5zHCNE – 28.6%; i5zHCNE – 88.9%; Fig. 7B,C, Supplemental Fig. S14, Table S14-15). The hindbrain expression domain was typically discrete, and not overlapping forebrain marker *otx2b:Venus* (Supplemental Fig. S15). This hindbrain expression driven by *HNF1B*/*hnf1ba* HCNEs is consistent with known expression of the endogenous genes in developing rhombomeres (Pouilhe et al. 2007; Choe et al. 2008). However, other sequences appear to be required for endoderm expression.

**Figure 7.**
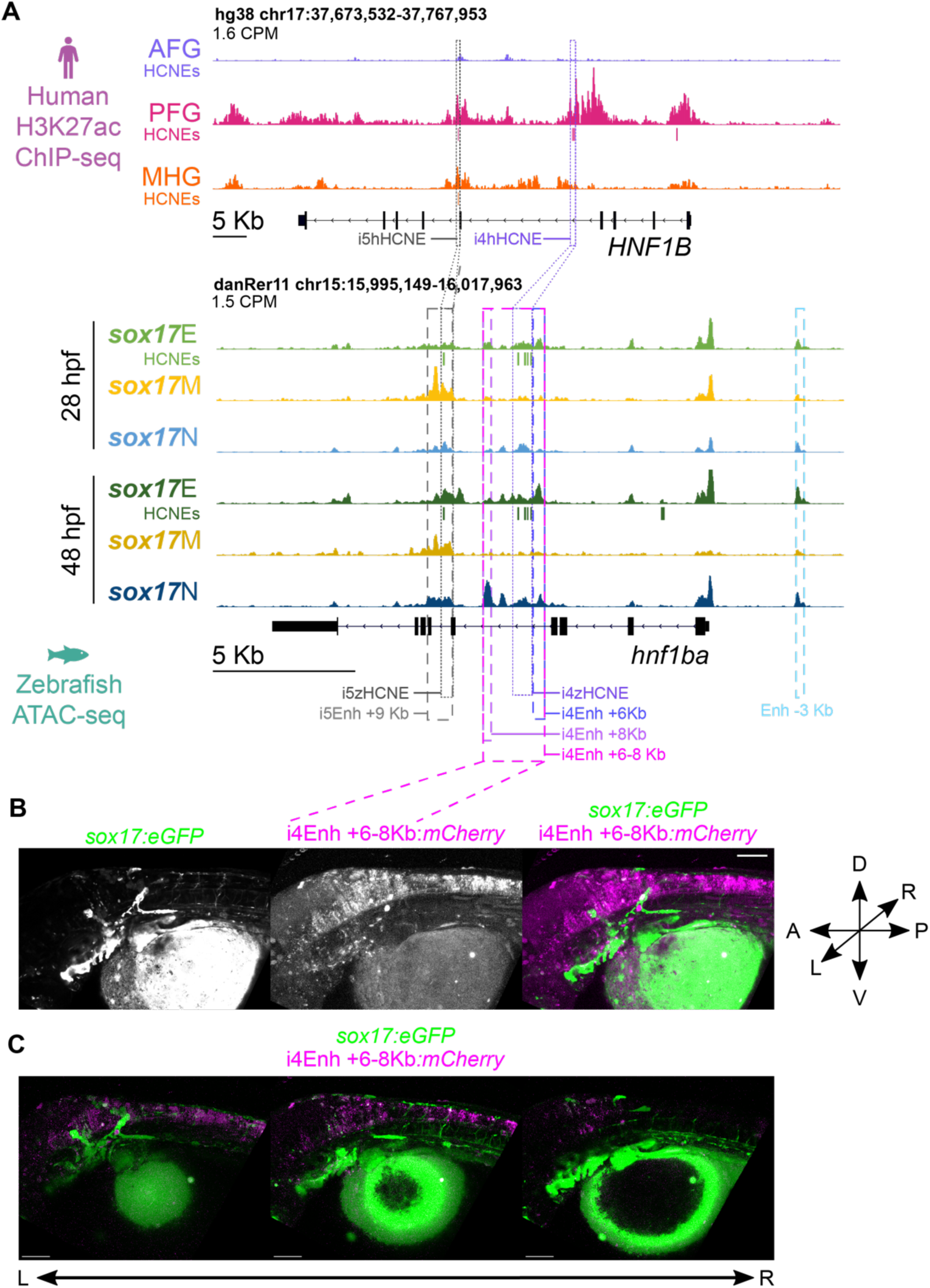
*HNF1B*/*hnf1ba* HCNE CRMs bearing the hallmarks of functionality in zebrafish endoderm and human PFG endoderm are sufficient to drive reporter expression in the developing hindbrain but not endoderm. (A) H3K27ac ChIP-seq and ATAC-seq data at human *HNF1B* and zebrafish *hnf1ba* loci respectively. HCNEs overlapping H3K27ac peaks and DARs are indicated in corresponding colours beneath tracks. Normalised peak heights in counts per million reads are indicated. HCNE and putative enhancer sequences cloned for reporter assays are outlined. (B) Lateral view of mCherry reporter expression driven by the i4Enh +6-8Kb enhancer at 48 hpf in *sox17:GFP* embryos injected with the reporter construct at the 1-cell stage. Maximum intensity z-projections of 27 z-slices from confocal images. (C) 3D rendering of confocal images showing 10 z-slices at a time going through the embryo along the z-axis from the left to right side. D = dorsal, R = right, P = posterior, V= ventral, L= left, A = anterior. Scale bars = 100 µm.

We note that *hnf1ba* intron 5 exhibits chromatin accessibility in the *sox17*M population, and both i5zHCNE and i5hHCNE consistently drive reporter expression in the zebrafish heart as well as the hindbrain (Fig. 7A, Supplemental Fig. S13, Table S15). We previously showed that *sox17:GFP* is co-expressed with *kdrl;mCherry* in the developing heart, consistent with known *sox17* expression in cardiac precursors in the lateral plate mesoderm (Chung et al. 2011; Johal et al. 2024). *Hnf1ba* expression has also been reported in the heart (Chi et al. 2008b). The i5HCNE enhancer therefore potentially regulates both cardiac and neural expression domains of *hnf1ba* but is insufficient for pancreas expression in spite of bearing the hallmarks of functionality in endoderm.

Overall we identify a subset of HCNEs bearing hallmarks of functionality in developing zebrafish and human endoderm during early organogenesis. These HCNEs are strongly enriched for TFBSs of factors critical during endodermal organ development and strongly associated with genes controlling craniofacial and pancreas development. However, the putative HCNE target genes governing pancreas development also have key roles in the developing nervous system, and the HCNE enhancers at the *HNF1b*/*hnf1ba* locus are sufficient to drive expression only the *HNF1b*/*hnf1ba* neural and cardiac but not endodermal expression domains. The present evidence suggests that the HCNEs are likely to be critically important for endoderm development, but must operate within a complex regulatory landscape to influence endodermal expression.

## Discussion

Our initial intention was to identify putative CRMs functioning in zebrafish endoderm during early organogenesis, before identifying those with clear homology with CRMs functioning in human endoderm. While our strategy to use transgenic reporter zebrafish to enrich for the developing endoderm was successful, we found that relatively few genomic regions are uniquely accessible in the endoderm. Rather, while many genomic regions exhibit enhanced accessibility in the *sox17*E population over the *sox17*M haematovascular cells, the majority of these regions are similarly accessible in the *sox17*N population. This suggests that any given putative endoderm CRM is likely to also be accessible in some other non-endodermal cell types. Consistent with this, amongst the enhancers identified in zebrafish and human endoderm, the HCNE enhancer in introns 4 of *HNF1B*/*hnf1ba* is capable of driving hindbrain expression, while the intron 5 enhancer drives both hindbrain and cardiac expression. In general, endoderm CRMs therefore appear unlikely to be endoderm-specific. Remarkably, despite being identified in endoderm, neither *HNF1B*/*hnf1ba* HCNE enhancer is individually capable of driving endodermal gene expression. This is most likely due to additional regulatory sequences being required to permit endodermal gene expression. Future analysis of *HNF1B*/*hnf1ba* regulation should therefore consider the combined actions of multiple CRMs, including analysis of whether the cognate *HNF1B*/*hnf1ba* promoters are required for the HCNE enhancers to influence endodermal gene expression. Analysis of chromatin accessibility and histone modifications at the single-cell level within developing tissues would also potentially allow identification of unique patterns of CRM accessibility indicative of combinatorial CRM action permitting target gene expression in specific cell types. Furthermore, it is possible that the identified *HNF1B*/*hnf1ba* HCNE CRMs may be necessary for *HNF1B*/*hnf1ba* endodermal expression even though they are not sufficient. Future studies should therefore involve production of CRISPR knockout zebrafish and/or human pluripotent stem cells followed by analysis of pancreas development.

### A common neuro-pancreatic gene regulatory and cis-regulatory programme?

Many similarities between endocrine cells of the pancreas and neurons have been noted over the years including production of polypeptide hormones, neurotransmitters and their receptors, and also common chromatin methylation signatures (Pearse and Polak 1971; Fujita et al. 1980; Gonoi et al. 1994; Maechler and Wollheim 1999; van Arensbergen et al. 2010). Furthermore, many of the TFs required for development of the endocrine pancreas have key roles in the developing nervous system throughout vertebrates, including *HNF1B*, *FOXA2*, *ISL1*, *PTF1A*, *ONECUT1*, *NEUROD1*, *HES1*, *PAX6*, *SOX4* and *MEIS2* (Ang and Rossant 1994; Ishibashi et al. 1995; Pfaff et al. 1996; Miyata et al. 1999; Sun and Hopkins 2001; Haubst et al. 2004; Hutchinson and Eisen 2006; Nolte et al. 2006; Hori et al. 2008; Potzner et al. 2010; Dal-Pra et al. 2011; Coolen et al. 2012; Roy et al. 2012; Machon et al. 2015; Jin and Xiang 2019). This is consistent with HCNE CRMs identified by our analyses of zebrafish and human endoderm being associated with both pancreas and nervous system development. Remarkably, TFBSs significantly enriched within these HCNE CRMs include those of TFs also operating during pancreas and nervous system development including *PTF1A*, *PDX1*, *FOXA2*, *ISL1*, *SOX4 NEUROD1* (Fig. 5). Given the *HNF1B*/*hnf1ba* endoderm HCNE CRMs drive expression in the developing nervous system, it is tempting to speculate that as well as there being a common set of TFs governing endocrine pancreas and nervous system development (neuro-pancreatic TFs), these TFs also act together within common CRMs that function in both the pancreas and nervous system. However, a broader analysis of HCNE CRMs performed at scale would be necessary to draw such conclusions more confidently. Since any given HCNE CRM is not necessarily individually sufficient to drive neural and pancreatic gene expression, future work will require both reporter assays and genetic deletion studies to test the effects on putative target genes. Combinatorial analysis of CRMs in both reporter assays and deletion studies will also likely be required to fully characterise the roles of HCNE CRMs regulating any given gene.

Common gene regulatory programmes spanning pancreas and nervous system development potentially implies co-option of an ancient gene regulatory programme from a common ancestral cell type. Indeed, all deuterostomes have a nervous system but only vertebrates have a proper pancreas. Thus, logically a common gene regulatory programme could have arisen during the evolution of the nervous system and subsequently been co-opted during pancreatic evolution. This is supported by analysis of sea urchin *Stongylocentrotus purpuratus* larvae, which identified a subset of neurons derived from cells with a putative “pre-pancreatic” signature consisting of expression of homologues of multiple neuro-pancreatic TFs including *SpSoxC* (homologous to *SOX4*) and *SpPtf1a* (Perillo et al. 2018). However, the endodermal HCNE CRMs identified are not clearly apparent in the lamprey genome. It therefore seems likely that though combined action of neuro-pancreatic TFs in the nervous system predates emergence of the vertebrate lineage, the HCNE CRMs they are predicted to bind are likely to have evolved after the divergence of jawless and jawed vertebrates.

### Endodermal HCNE CRMs and complex vertebrate-specific patterning

Our analyses suggest that endodermal HCNE CRMs may participate in patterning anterior structures including the craniofacial skeleton and ear (Fig. 4H, Supplemental Figs S9,11,12). While the anterior endoderm does not contribute directly to the craniofacial skeleton, it is considered to have a key role as a conserved signalling centre governing the patterning of anterior neural crest derivatives. Indeed, disruption of multiple signalling pathways within or emanating from the endoderm including Hedgehog, FGF and BMP, as well as their upstream regulators leads to craniofacial defects (reviewed in (Graham et al. 2005; Grevellec and Tucker 2010). Furthermore, the anterior endoderm has multiple roles during ear development, contributing directly to the eardrum and Eustachian tube but also acting as a signalling centre patterning the broader ear (Grevellec and Tucker 2010; Anthwal and Thompson 2016). Indeed, defects in pharyngeal endodermal gene expression have been suggested to lead to altered patterning of anterior structures in human syndromes. For example, DiGeorge syndrome is characterised by defects in the pharyngeal apparatus and is principally considered to be caused by *TBX1* haploinsufficiency. Tissue-specific *Tbx1* knockout in the pharyngeal endoderm in mouse leads to multiple defects in none-endodermal tissues including cleft palate and absent outer and middle ear (Arnold et al. 2006). It therefore seems likely that endodermal HCNE CRMs participate in anterior endoderm functions governing craniofacial patterning. However, there are likely to be key commonalities between the zebrafish lateral line system and mechanosensory cells of the mammalian inner ear (Duncan and Fritzsch 2012). Since anterior lateral line neurons are likely to be present in the *sox17*E population it is possible that some HCNEs contributing to ear-related terms are accessible in lateral line neurons rather than the endoderm. However, such HCNEs are still within human endoderm H3K27ac peaks. Nevertheless, our analyses reveal a set of HCNE CRMs bearing hallmarks of functionality in both human and zebrafish that should be prioritised for analyses of gene regulatory control of craniofacial and ear development.

### Individual HCNE CRMs appear to have a distinct regulatory logic

Highly conserved CRMs have been suggested to be constrained by the action of TF complexes requiring consistent configuration of TFBSs (Guturu et al. 2013; Long et al. 2016; Jindal and Farley 2021). Our analysis of PFG HCNE CRMs suggests highly conserved patterns of TFBS co-occurrence (Fig. 6). However, relatively few TF pairs show significant enrichment for specific TF spacing across different HCNEs within the same species. This suggests that either the selective pressure maintaining HCNEs is not rigid sequence requirements for TF complex binding, or that TF complexes requiring such rigid grammar do not act widely across HCNE CRMs. It is also possible that depending on the configuration DNA binding domains within TF complexes, they are robust to variability in TFBS spacing. An alternative model to explain conserved patterns of TFBS co-occurrence without rigid spacing relates to recently proposed “dependency grammar” (Jindal and Farley 2021). This considers that multiple parameters including TFBS affinity, order, spacing and orientation are tuned by selective pressures to yield viable patterns and levels of expression from key enhancers. Thus, interplay between spacing and affinity of sites dictates the functional capabilities of enhancers without either parameter needing to be rigidly set (Farley et al. 2015; Farley et al. 2016). This may be linked to suboptimisation of enhancers via hinderance of combined TF binding, preventing overactivation (Farley et al. 2015). This is especially important for *trans-dev* genes, where their overactivation or ectopic expression may lead to harmful alterations in cell fate assignment during development. Enhancer configuration with genuine but less rigid grammatical logic may also offer an evolutionary advantage since such enhancer are more likely to be robust to minor sequence changes without dramatic alterations in function.

Given our results suggest combinatorial rules in terms of TFBS co-occurrence without rigid spacing in PFG HCNE CRMs, this supports the dependency grammar model where co-regulated genes are regulated by similar sets of TFs, but the tuning of grammatical parameters varies to ensure correct functional outputs. This tuning via dependency grammar is likely to be critical, hence the high degree of conservation of the HCNEs.

Overall our analyses reveal a compendium of HCNE CRMs acting across the 400 million years of evolution separating zebrafish and humans. These HCNE CRMs should be prioritised for analysis of commonalities of gene regulatory control programmes controlling neural and pancreatic development, and also the role of endoderm in patterning the craniofacial apparatus. Our analyses reveal consistent patterns of TFBS co-occurrence in endoderm HCNE CRMs that should form the basis of future studies of TF combinatorial and competitive binding to reveal conserved mechanisms tuning the correct spatiotemporal expression of *trans-dev* genes.

## Methods

### Zebrafish strains and transgenics

ABix, *Tg(-5.0sox17:GFP)^ha01Tg^* (Mizoguchi et al. 2008), *Tg(kdrl-HsHRAS:mCherry)^s896^* (Chi et al. 2008a) and *Tg(gata1a:DsRed*)*^sd2Tg^* (Traver et al. 2003) fish were reared as described (Westerfield 2007). All zebrafish studies fully complied with the UK Animals (Scientific Procedures) Act 1986 as implemented by University of Warwick.

### Preparation of cells for RNA-seq and ATAC-seq

Embryos were dechorionated using 1 mg/ml pronase and deyolked in calcium free Ringer solution (0.5 M EDTA, 116 mM NaCl, 2.9 mM KCl, 5 mM HEPES). Deyolked embryos were dissociated in 20 mg/ml Collagenase (Sigma #C8176-25MG), 0.05% Trypsin with EDTA (Gibco #15400-054) in 1x Hank’s Balances Salt Solution (HBSS, Gibco #14185-045) and homogenised using a pipette tip. The reaction was stopped by addition of foetal bovine serum to 5%, followed by dilution in HBSS+/+ (1X HBSS, 0.25% (w/v) BSA, 1 mM HEPES). Cells were pelleted and washed in HBSS +/+ buffer followed by filtration through a 70 μm cell strainer (Millex-GP). Cells were centrifuged for 5 minutes at 750 xg and re-suspended in HBSS+/+ buffer and transferred to polypropylene tubes pre-coated with 5% FBS in 1x PBS. Cell sorting was performed using a Becton Dickinson FACSAria Fusion Cell Sorter with a 100 μm nozzle and sheath fluid pressure of 25 pounds per square inch (psi). Debris was eliminated based on side scatter area (SSC-A) and forward scatter area (FSC-A) and singlet cells selected based on forward scatter area (FSC-A) and forward scatter width (FSC-W). Next the single cells were sorted based on fluorescence signal from green fluorescent protein *EGFP* (B488-530) and red fluorescent proteins *mCherry* and *DsRed* (YG561-610). Flow cytometry was operated and analysed using BD FACSDiva^TM^ software.

### RNA-seq

Total RNA was prepared from sorted cells using QIAGEN RNeasy Mini kit with on-column DNase I treatment (QIAGEN) following the manufacturer’s protocol. 10 ng total RNA per sample was use to construct sequencing libraries using the NEBNext Ultra II Directional RNA Library Prep kit for Illumina according to the manufacturer’s instructions and sequenced using Novaseq 6000-S4-type flow cell. Reads were mapped onto zebrafish genome, danRer11 using STAR (Dobin et al. 2013). HTSeq was used to count aligned reads per gene (Anders et al. 2015). The raw count matrix was imported into iDEP (Ge et al. 2018), and differentially expressed genes (DEGs) identified using default parameters. Heatmaps of DEGs were produced using Morpheus (https://software.broadinstitute.org/morpheus).

### ATAC-seq

50,000 sorted cells per sample were used for OMNI-ATAC-seq using adapted methods outlined by (Buenrostro et al. 2015) and (Corces et al. 2017). All steps were conducted on ice or in centrifuges at 4°C. Cells were washed in ice cold 1x PBS prior to centrifugation at 500 xg for 5 minutes. Cell pellets were resuspended with cell lysis buffer (10mM Tris-HCl pH 7.5, 10mM NaCl, 3mM MgCl_2_, 0.1% (v/v) NP-40, 0.1% (v/v) Tween-20, 0.01% (v/v) Digitonin) and were spun down at 4°C 500 xg for 10 minutes. Pelleted nuclei were resuspended in transposition reaction mix (1X Tagment D buffer (Illumina #15027866), 5% (v/v) Tagment DNA Enzyme 1 (Nextera), PBS, 0.1% (v/v) Tween-20, 0.01% (v/v) Digitonin). Transposition reaction was incubated shaking at 1,000 rpm at 37°C for 30 minutes. DNA was isolated using Qiagen MinElute PCR Purification Kit and eluted using 22μl elution buffer. The remaining ATAC-seq library preparation was performed as described previously (Buenrostro et al. 2015). Bioanalyzer High Sensitivity DNA Analysis kit (Agilent) was used to test the quality of the purified libraries and ensure fragment size periodicity with intervals of around 200 bp. Bioanalyzer results were used to estimate concentrations and inform dilutions to be conducted before sending samples for sequencing. Libraries were sequenced by the Genomics Facility at the University of Warwick with Illumina NextSeq 500 using the High Output Kit v2.5 (FC-404-2002), and by Novogene (UK) Company Limited using Novaseq 6000-S4-type flow cell. Each sample used different barcoded reverse primers, allowing for the samples to be multiplexed for sequencing.

### ATAC-seq data analysis

Reads were trimmed from the 3’ end to a uniform 75 bp using seqtk so that mapping of reads was not impacted by sequencing read length. Adapter sequences were trimmed using Trimmomatic version 0.39 (Bolger et al. 2014). Data quality control and confirmation of adapter removal was performed using FastQC (Andrews 2010). Reads were aligned to the genome reference consortium zebrafish build 11 (GRCz11v97) using Bowtie2 (Langmead and Salzberg 2012) using “Very sensitive” and 2 kb maximum fragment length for paired-end alignments. Samtools (Li et al. 2009) was also used to filter out reads that mapped to the mitochondrial genome, duplicated reads and low-quality reads with a MAPQ score < 22.

Accessible regions (peaks) were called using MACS2 (Zhang et al. 2008; Gaspar 2018) with default parameters except: broad peak setting, mappable genome size of 1.4e9, false discovery rate cut-off of 0.05, bandwidth of 300bp, and a high-confidence fold-enrichment between 5 and 50. To control for Tn5 sequence-specific signature bias, peaks were called against an ATAC-seq control library produced from purified genomic DNA (Buenrostro et al. 2013).

Differentially accessible regions (DARs) between the cell populations and time points, were identified in R version 4.2.1 (R Core Team, 2021) using DiffBind version 3.0.15 (Buenrostro et al. 2013) with the DESeq2 option (Love et al. 2014), on the broad peaks files (.broadPeak) produced by MACS2. Summits were set to false so that the summit heights (read pileup) and locations were not calculated for each peak, allowing the whole peak size to be considered. DARs identified with FDR <= 0.01 DARs were used for subsequent analysis. DARs were assigned to the nearest TSS and distributions of DARs relative to genes assessed using ChIPseeker 1.32.1 (Yu et al. 2015) and GRCz11 (danRer11) Ensembl gene annotation version 97.

Gene ontology (GO) term enrichment for biological processes, molecular functions and pathways was performed using over-representation analysis (ORA) in WebGestalt 2019 (Liao et al. 2019). FishEnrichr (Chen et al. 2013; Kuleshov et al. 2016) was used to identify zebrafish anatomical structures associates with genes lists.

Integrated genome viewer (IGV) version 2.5.0 (Robinson et al. 2011) was used to visualise mapped ATAC-seq and ChIP-seq reads, DARs and HCNEs. ATAC-seq peaks from biological replicates were merged using Samtools (Li et al. 2009) to show representative peaks. Tracks were normalised using the CPM function in deepTools bamCoverage (Ramirez et al. 2016).

To produce heatmaps of ATAC-seq data BAM files were downsampled to equalize read numbers between conditions using Picard. Heatmaps were produced using seqMINER v1.3.4 using the KMeans enrichment linear clustering normalization method (Ye et al. 2011).

### ChIP-seq data analysis

Previously published H3K27ac ChIP-seq data was downloaded from GEO: Series GSE52658 (Loh et al. 2014). Reads were aligned to hg38 using Bowtie2 (Langmead and Salzberg 2012) with default parameters apart from-N 1. ChIP-seq peaks were called using MACS2 (Zhang et al. 2008; Gaspar 2018) using default settings against cognate controls. A more stringent q-value of <10^-8^, instead of the default <0.05, was qualitatively determined as an appropriate cut off for the called peaks by looking at the peaks around key genes known to be expressed in endodermal cell populations, as well as genes expressed specifically in non-endodermal cell populations.

### Analysis of Highly Conserved Non-coding Elements (HCNEs) overlapping ATAC-seq and ChIP-seq datasets

HCNEs comparing human (hg38) and zebrafish (danRer10) genome builds using a window size of 30 bp and 70-100% percent identity threshold were downloaded from ANCORA (http://ancora.genereg.net) (Engstrom et al. 2008). danRer10/hg38 HCNEs were converted to danRer11/hg38 using liftOver (Hinrichs et al. 2006). Bedtools intersect (Quinlan and Hall 2010) was used to determine if danRer11/hg38 HCNEs overlap with zebrafish danRer11 DARs from ATAC-seq and human hg38 ChIP-seq peaks.

Functional and anatomical enrichment analyses were performed for hg38 HCNE sets using Genomic Regions Enrichment of Annotations Tool (GREAT) version 4.0.4 with default settings (McLean et al. 2010; Tanigawa et al. 2022). Dual analyses were performed with background regions set to whole genome or a list of all HCNEs. Heatmaps of the top 20 enriched terms per set of HCNEs were selected based on hypergeometric false discovery rate corrected q-values (HyperFdrQ).

### Motif enrichment analysis

danRer11 and hg38 HCNE genomic sequences were extracted using BED2FASTA in the MEME Suite (Bailey et al. 2015), followed by *de novo* and known motif enrichment analysis using HOMER v4.11.1 (Heinz et al. 2010). For known motif enrichment analysis the top 30 most significant motifs in each HCNE set (P < 1 x 10^-8^) were considered leading to a combined set of 60 motifs. A heatmap of indicating enrichment of these motifs in each HCNE set was produced and clustered using Morpheus (https://software.broadinstitute.org/morpheus). To annotate expression of transcription factors corresponding to these motifs RPKM values from relevant human endoderm RNA-seq datasets were downloaded from GEO: Series GSE52658 (Anterior Foregut, Posterior Foregut and Mid/Hindgut endoderm) (Loh et al. 2014) and GSE216266 (passage 0 pancreatic progenitors) (Jarc et al. 2024). Cooccurrence of selected motifs within HCNEs were analysed using Paired Motif Enrichment Tool (PMET implemented at https://pmet.online and https://github.com/duocang/PMET-Shiny-App) (Rich-Griffin et al. 2020) on genomic intervals using default parameters. PMET output was used to produce a matrix using the acast function in R, prior to producing a heatmap using Conditional Formatting in Microsoft Excel. Pairwise analysis of spacing of motifs from HOMER was performed using SpaMO in MEME Suite (Whitington et al. 2011; Bailey et al. 2015).

### Cloning of reporter constructs for reporter assays

Putative enhancer and HCNE elements of interest were PCR amplified from zebrafish or human genomic DNA using custom primers containing *att*B4 and *att*B1 sequences and Q5 ® High-Fidelity DNA Polymerase (NEB) and cloned by Gateway recombination into pDONR-P4-P1R (Thermo Fisher Scientific). To allow visualization of reporter activity in *Tg(sox17:GFP)* zebrafish, constructs were generated to express mCherry downstream of the putative enhancer. To do this published plasmids pENTRbasEGFP (Addgene #22453), pENTREGFP2 (Addgene #22450) and pDESTtol2pACrymCherry (Addgene #64023) (Villefranc et al. 2007; Berger and Currie 2013) were modified using NEBuilder HiFi DNA Assembly Master Mix (NEB) to switch *EGFP and mCherry* between the plasmids. To all use of the E1b promoter as the basal promoter in reporter assays the E1b promoter was amplified from *gata2a*-i4-E1b-GFP-Tol2 (kindly gifted by Dr Rui Monteiro) and inserted into pENTR *mCherry* by HiFi cloning. All primers used are listed in Supplemental Table S16). Following sequence verification, *hnf1ba* putative enhancer/HCNE, pENTR E1bP:*mCherry*, p3E-mcs1 (Addgene #49004) (Moore et al. 2013) and pDESTtol2pACry*EGFP* constructs were recombined by Gateway recombination (Thermo Fisher Scientific). This generated plasmids with putative *hnf1ba* enhancer elements upstream of the E1b promoter and mCherry, and the *cryaa* promoter upstream of EGFP.

### Embryo compound treatments, reporter assays and imaging

Embryos for imaging were incubated from 22 hpf with 0.003% PTU (1-phenyl 2-thiourea) to block pigmentation and improve optical transparency. Embryos were immobilised for imaging using 30 μg/ml tricaine (ethyl 3-aminobenzoate methanesulfonate E10521-10G MERCK) and mounted and orientated in 1% agarose moulds created using a 3D-printed stamp (Kleinhans and Lecaudey, 2019). Embryos were imaged using a Nikon ECLIPSE Ni, Zeiss Axio Zoom.V16 or Zeiss 980 confocal microscope. Images were analysed and processed using ImageJ/Fiji (Schindelin et al. 2012) version 2.9.0/1.52t. Processing included: adjusting the brightness and contrast setting, adding scale bars, creating and saving composite images, changing LUTs, and applying math log function to help deal with saturation. Bio-Formats Plugins for ImageJ (Linkert et al. 2010) was used to import files. Temporal-Colour Code was used to generate a 2D image from a stack of widefield fluorescent images, with each focal slice colour coded based on the Ice LUT. 3Dscript (Schmid et al. 2019), an ImageJ/Fiji plugin, was used to render 3D images and videos from confocal images. Independent transparency was used as the rendering algorithm.

To analyse the effects of retinoic acid (RA) signalling, embryos were treated from 5 hpf with either RA (Sigma-Aldrich R2625), diethylaminobenzaldehyde (DEAB Sigma-Aldrich D86256), BMS493 (APExBIO B7415) or vehicle (DMSO - 0.1%) as a control. Following treatment embryos were incubated in the dark to prevent degradation of the light sensitive chemicals.

For reporter assays, *Tg(sox17:GFP)* embryos were microinjected at the one cell stage with 25 pg reporter constructs combined with 25 pg Tol2 transposase capped mRNA produced from pCS2-Tol2. Embryos were screened using widefield fluorescent microscopes for expression of *cry:EGFP* in the lens at 48 hpf for confirmation of reporter construct integration into the genome. The location of *mCherry* expression in these embryos was observed using widefield fluorescent microscopes and scored. Embryos of interest were imaged further using confocal microscopes.

## Data access

All raw and processed sequencing data generated in this study have been submitted to the NCBI Gene Expression Omnibus (GEO; https://www.ncbi.nlm.nih.gov/geo/) under accession number GSEXXXXXX.

## Acknowledgements

This research was also funded, in-part, by the Wellcome Trust through a Wellcome Seed Award in Science (210177/Z/18/Z) and an MRC New Investigator Research Grant (MR/S021531/1) to ACN. DMF and RE were funded by the MRC Doctoral Training Partnership in Interdisciplinary Biomedical Research (MR/N014294/1). MW was funded in-part by the Quantitative Biomedicine Programme through Warwick’s Wellcome Institutional Strategic Support Fund (ISSF) award with match funding provided by the University of Warwick. SJ was funded by BBSRC Midlands Integrative Biosciences Training Partnership (BB/M01116X/1). We thank Rui Monteiro for kindly gifting us the *Tg(kdrl:mCherry)* and *Tg(gata1a:dsRed)* fish used in this study, Fiona Wardle for *Tg(sox17:GFP)* and Michael Smutny for *Tg(otx2b:Venus)*. pENTRbasEGFP, 599 p3E-mcs1 and pENTREGFP2 were gifts from Nathan Lawson (Addgene plasmid # 22453; http://n2t.net/addgene:22453;RRID:Addgene_22453;RRID:Addgene_22453;plasmid#49004;http://n2t.net/addgene:49004;RRID:Addgene_49004;Addgeneplasmid#22450;http://n2t.net/addgene:22450; RRID:Addgene_22450). pDESTtol2pACrymCherry was a gift from Joachim Berger & Peter Currie (Addgene plasmid # 64023; http://n2t.net/addgene:64023; RRID:Addgene_64023). We thank Fiona Wardle and Karuna Sampath for reagents. We thank Karuna Sampath, Andrea Zaucker and Andre Pires da Silva for valuable discussions, and Karuna Sampath, Jonathan Millar and Andre Pires da Silva for generous access to equipment. We also thank the University of Warwick Research Technology Platform (RTP) Aquatics facility for zebrafish care and, Ian Hands-Portman from the Warwick School of Life Sciences for imaging advice and support. We thank Warwick Integrative Synthetic Biology Centre (WISB) for flow cytometry access, and Sarah Bennett for initial training. WISB is a BBSRC/EPSRC Synthetic Biology Research Centre (BB/M017982/1) funded under the UK Research Councils’ Synthetic Biology for Growth programme. We thank the Warwick School of Life Sciences Genomics Facility for sequencing and technical support, and Kate Woolley-Allen, Pavle Vrljicak, Julia Lipecki, Paul Brown and Warwick Bioinformatics RTP for bioinformatics advice. We thank Xuesong Wang for support with the use of PMET.

## Author contributions

A.C.N. conceived the study. A.C.N., S.O. and T.B. supervised the study.

A.C.N. and D.M.F. designed the experiments. D.M.F., R.E., M.W., S.J., Y.L. and A.C.N. performed all the experiments. D.M.F., R.E., M.W., S.J. and A.C.N. performed data analysis, and visualization. A.C.N. and D.M.F. wrote the original draft. All authors read and approved the final manuscript.

