## Supplemental Figures and Tables 2,3,11 and 13-16 for "Functional genomics analysis of developing zebrafish and human endoderm reveals highly conserved *cis*-regulatory modules controlling vertebrate organogenesis"

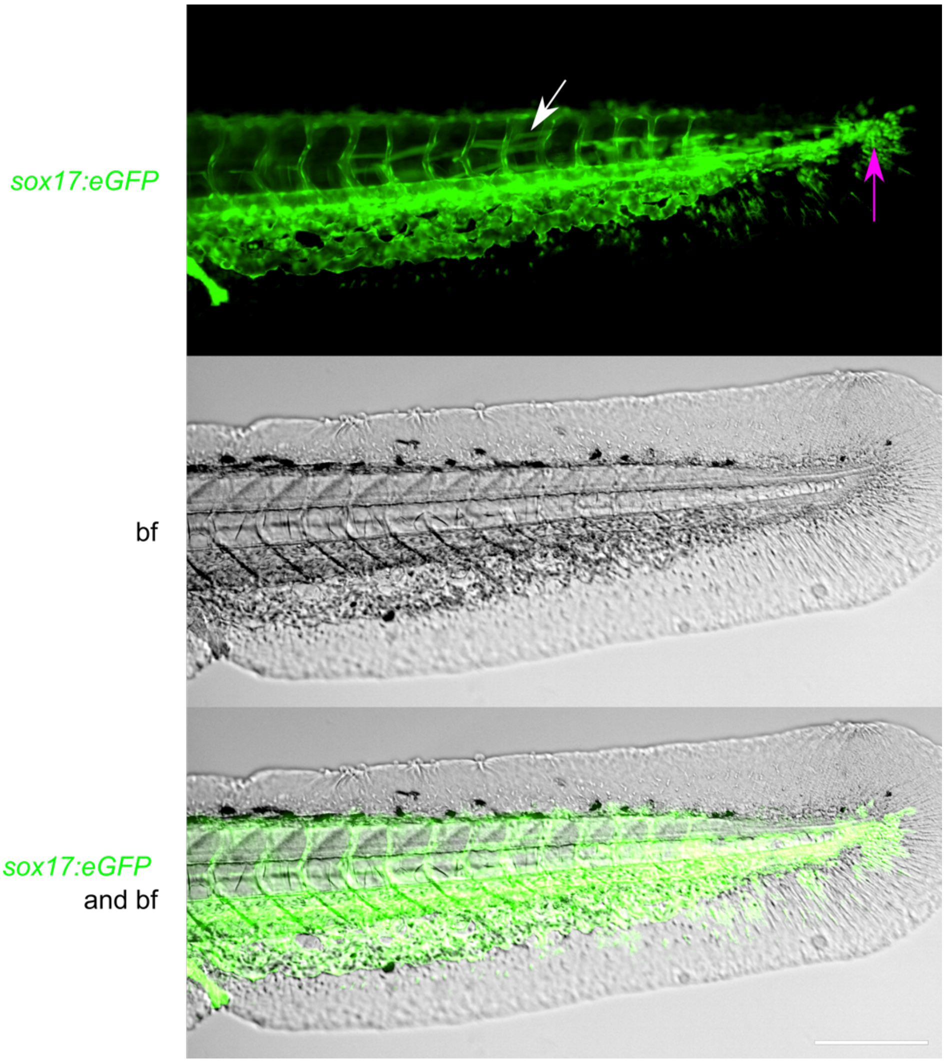


***Supplemental Figure 1: Tg(sox17:EGFP) 48 hpf embryos show expression in the median fin fold, and muscle cells in the tail***

*Images of lateral orientated 48 hpf embryos. Math Log function has been applied to the images. White arrow is pointing at muscle cells that are eGFP positive. Pink arrow points at eGFP in the median fin fold cells. White scale bar in composite image represents 500mm.*

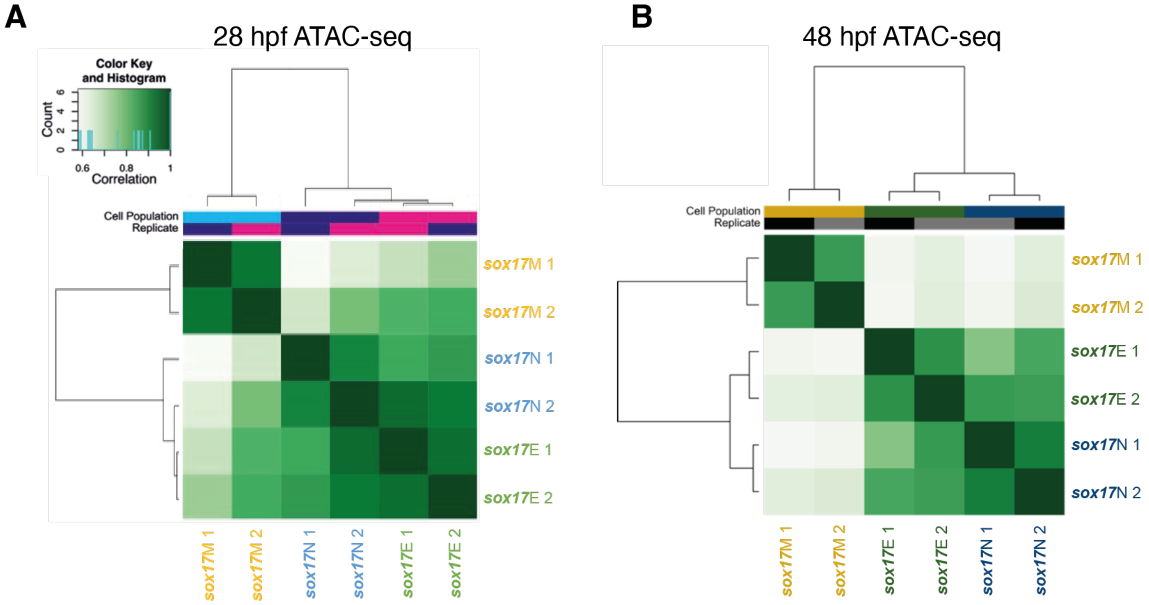


***Supplemental Figure 2. Biological replicates of sorted populations cluster together***
*Pearson correlation heatmap based on MACS2 scores for ATAC-seq data at 28 hpf (A) and 48 hpf (B).*

***Supplemental Table 2: Number of DARs between sorted cell populations at 28 and 48 hpf***

| **Cell populations compared** | **Comparison type** | **Number of regions (DARs) exhibiting greater accessibility per timepoint** | | | |
|---|---|---|---|---|---|
| | | 28 hpf | | 48 hpf | |
| | | FDR<0.05 | FDR <0.01 | FDR <0.05 | FDR <0.01 |
| Sox17M vs Sox17E | Sox17M> Sox17E | 10,300 | 6,267 | 7,823 | 5,500 |
| | **Sox17E> Sox17M** | **23,344** | **12,898** | **46,703** | **28,172** |
| Sox17N vs Sox17E | Sox17N> Sox17E | 1,102 | 107 | 6,209 | 1,557 |
| | Sox17E> Sox17N | 599 | 91 | 5,692 | 2,542 |
| Sox17N vs Sox17M | Sox17N> Sox17M | 29,133 | 19,557 | 60,282 | 41,719 |
| | Sox17M> Sox17N | 12,367 | 8,521 | 8,741 | 5,293 |

***Supplemental Table 3: Number of DARs between 28 and 48 hpf in the sorted cell populations***

| **Cell populations compared** | **Comparison type** | **Number of regions (DARs) exhibiting greater accessibility** | |
|---|---|---|---|
| | | FDR <0.05 | FDR <0.01 |
| **Sox17E** | 28 hpf > 48 hpf | 985 | 395 |
| | 48 hpf > 28 hpf | 1,707 | 22 |
| **Sox17M** | 28 hpf > 48 hpf | 4,828 | 2,317 |
| | 48 hpf > 28 hpf | 1,090 | 478 |
| **Sox17N** | 28 hpf > 48 hpf | 5,788 | 3,100 |
| | 48 hpf > 28 hpf | 4,136 | 1,264 |

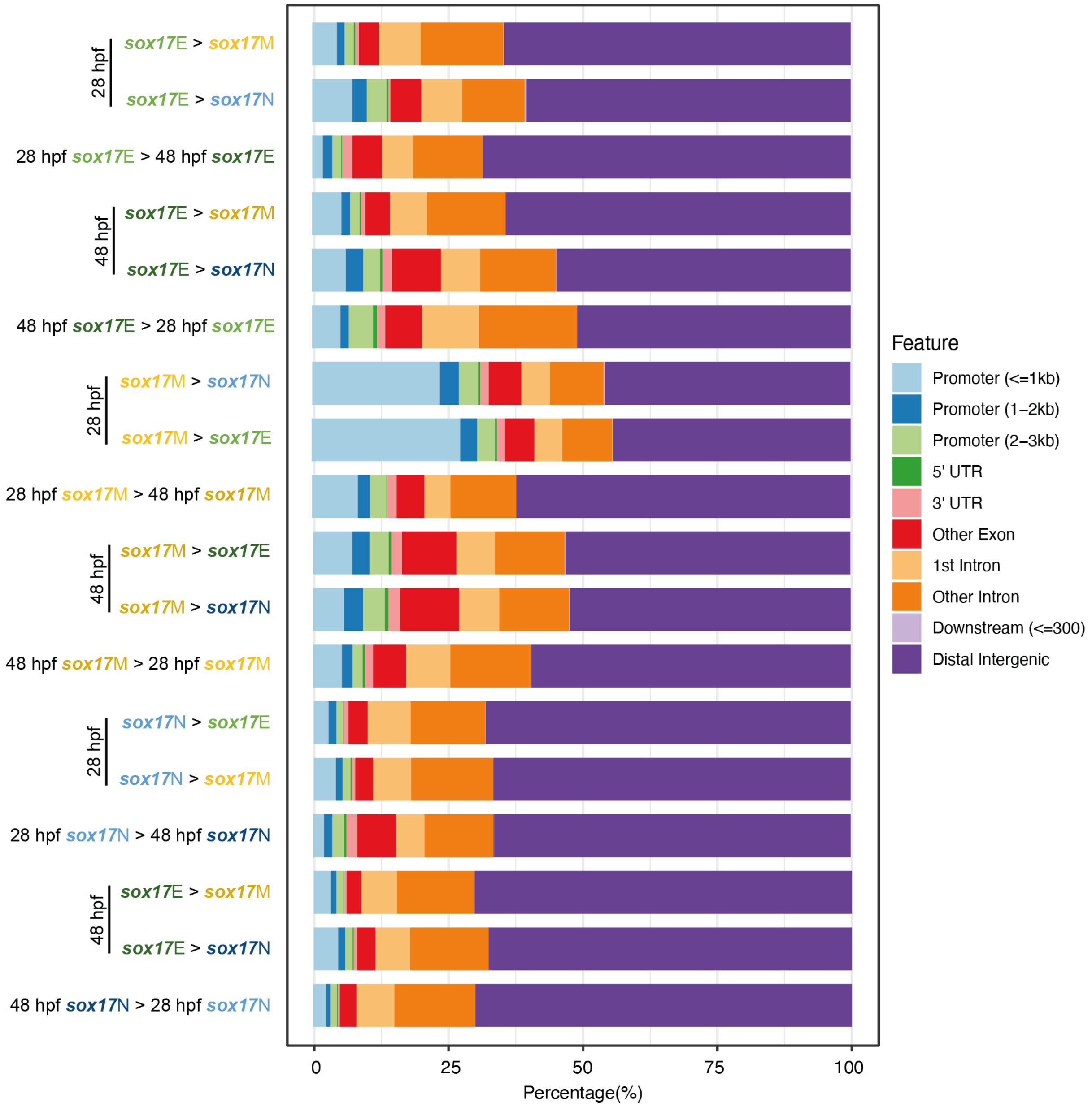


***Supplemental Figure 3. Genomic distribution of DARs relative to gene annotations***

*DARs from all pairwise comparisons were related to zebrafish danRer11 gene annotations using ChIPseeker. The percentage DARs in each location bin are indicated.*

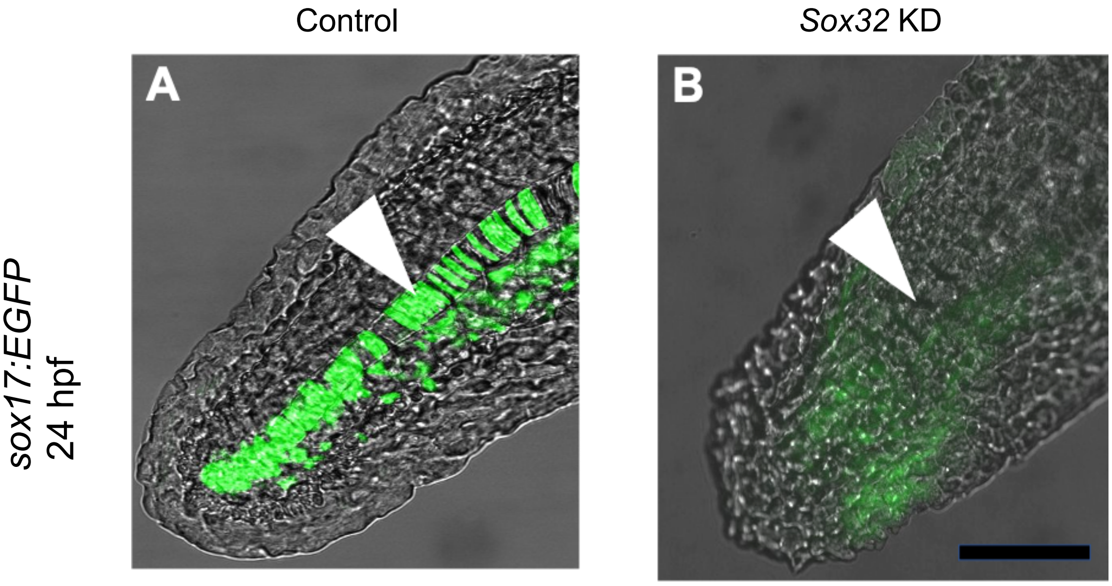


***Supplemental Figure 4. GFP+ cells in the posterior notochord of sox17:EGFP arise from sox32-dependent cells.*** *Sox17:EGFP embryos injected at the one-cell stage with either control morpholino (A) or sox32 morpholino (B) and imaged at 24 hpf. The posterior notochord is shown as an overlay of fluorescence and brightfield images. Arrowhead indicates posterior notochord. Scale bar, 0.25 mm.*

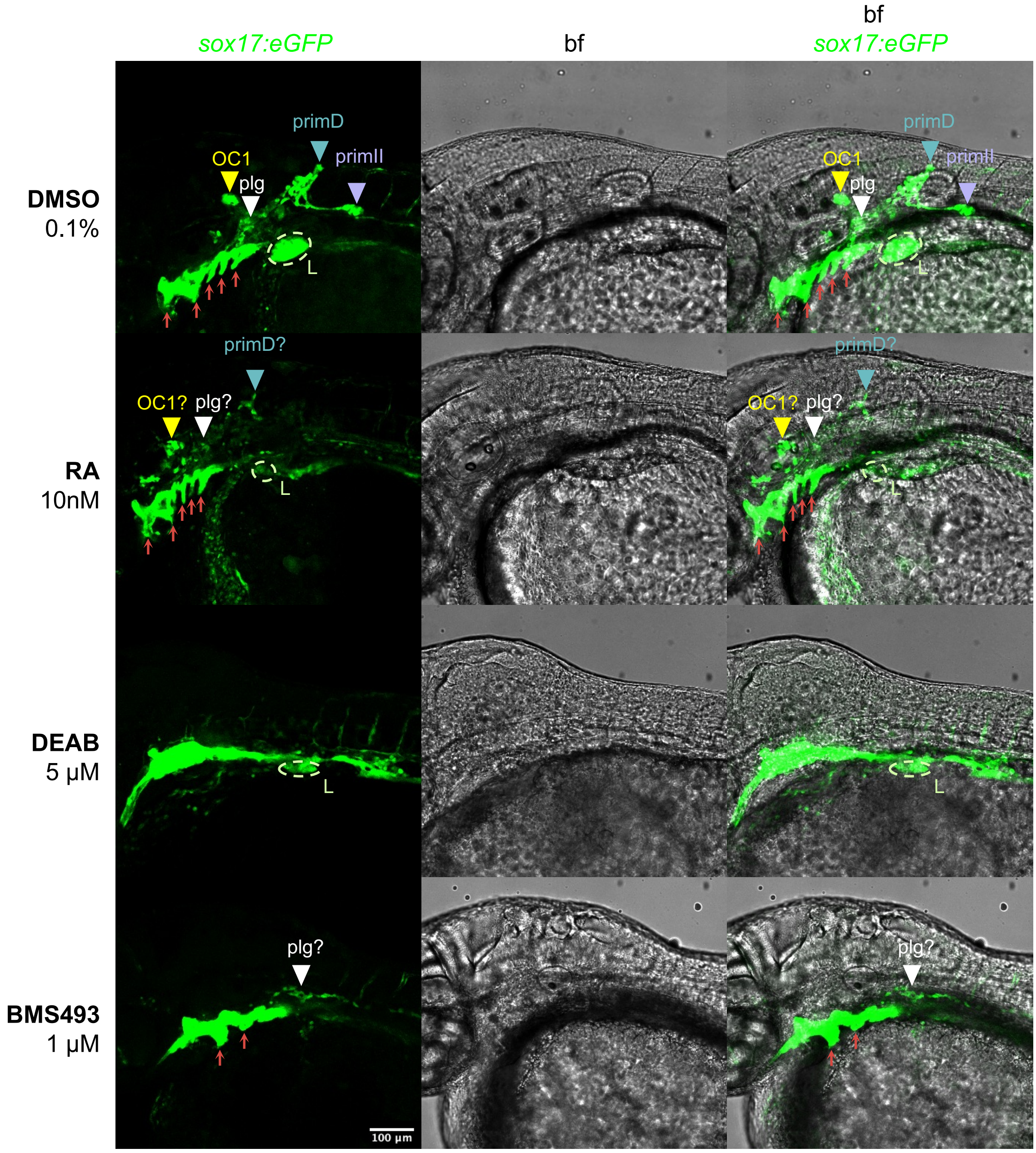


***Supplemental Figure 5. sox17:EGFP expression in the posterior lateral line is sensitive to RA treatment***

*Z-projections of 17 slices of confocal images of 48 hpf Tg(sox17:EGFP) embryos treated with DMSO (0.1%), RA (10nM), DEAB (5mM) or BMS493(1mM). Laterally orientated. Images from left to right are firstly sox17:EGFP, secondly bf and thirdly overlay of bf and eGFP (green). Dorsal is top, posterior is right, ventral is bottom, anterior is left. Scale bar = 100mm. Key estimated expression domains are marked and labelled as: OC1 = organ of Corti 1 (yellow arrow head), plg = posterior lateral line ganglion (white arrow head), primD = dorsal primordium (teal arrow head), primII = second primordium (purple arrow head), L = liver (light green dotted line), pharyngeal pouches = pp (red arrow heads). Question marks denote regions that are expected to be the expression domain based on morphology and location.*

***Supplemental Table 11. Number of H3K27ac peaks called in each cell population drops with more stringent q-values***

| Cell population | Number of peaks q-value <0.05 | q-value $<10^{-8}$ |
|---|---|---|
| AFG | 153,956 | 69,853 |
| PFG | 150,374 | 72,194 |
| MHG | 129,060 | 56,159 |

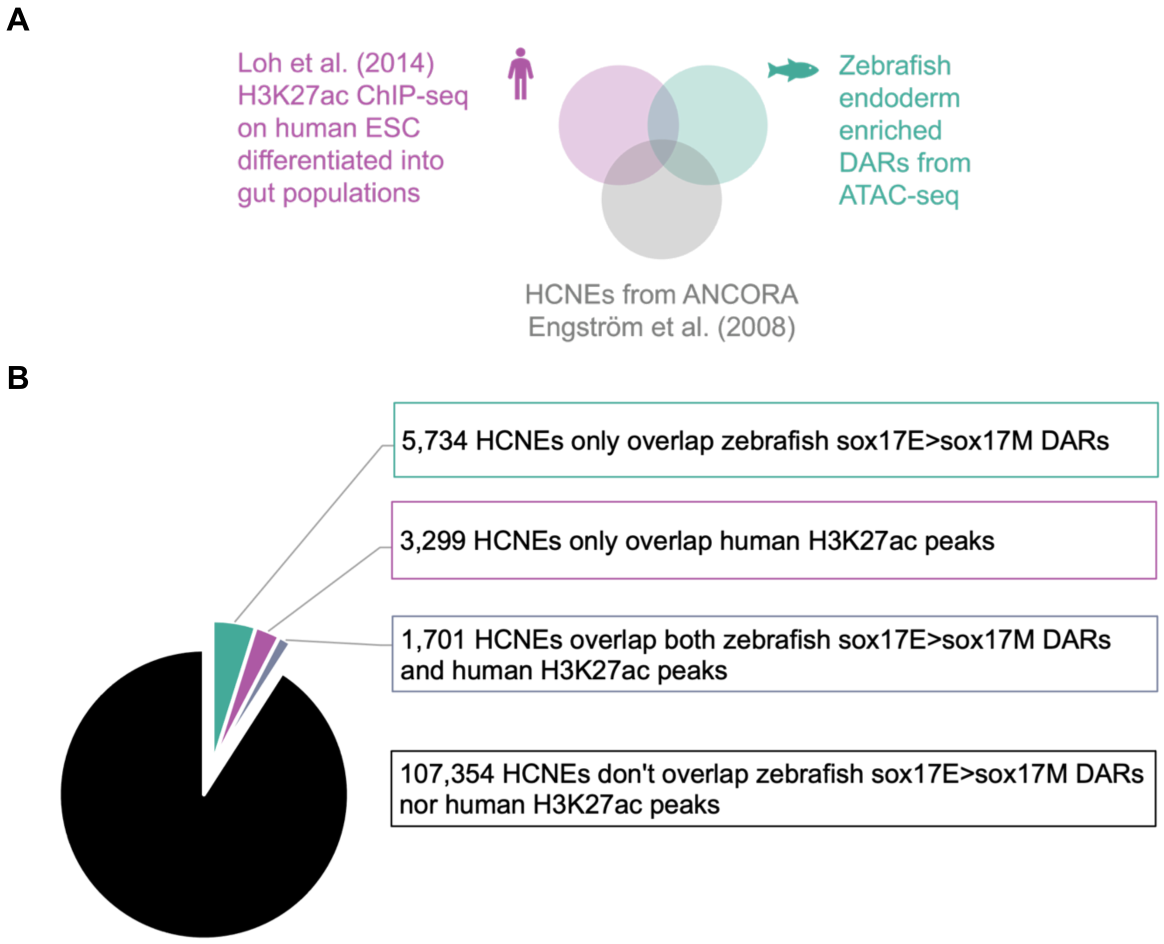


***Supplemental Figure 6. Human endoderm H3K27ac peaks and/or zebrafish sox17E>sox17M DARs overlap with 9% of zebrafish/human HCNEs***
***A.*** *Schematic of the overlap of HCNEs from ANCORA with the functional genomics data of H3K27ac peaks in hESCs differentiated into anterior posterior patterned endodermal cells from Loh et al. (2014) and the endoderm enriched sox17E>sox17M DARs from 28 and 48 hpf zebrafish embryos collected by us using ATAC-seq.* ***B.*** *Pie chart showing the number of HCNEs overlapping the two data sets aforementioned.*

***Supplemental Table 13. Number of HCNEs overlapping zebrafish sox17E>sox17M DARs and H3K27ac peaks in anterior-posterior patterned endoderm cell populations derived from hESCs***

| | | What HCNEs overlap | | | | | Number of HCNEs | | |
|---|---|---|---|---|---|---|---|---|---|
| | | Zebrafish sox17E>sox17M DARs | | Human H3K27ac peaks | | | in overlap | in category | total |
| **Name of category** | **Name of overlap** | **28 hpf** | **48 hpf** | **AFG** | **PFG** | **MHG** | | | |
| **Only overlap sox17E>sox17M DARs** | 28 | | | | | | 2,392 | | |
| | 48 | | | | | | 1,886 | | |
| | 28 48 | | | | | | 1,456 | | 5,734 |
| **Overlap both sox17E>sox17M DARs and H3K27ac peaks** | 28 AFG | | | | | | 104 | | |
| | 28 AFG PFG | | | | | | 169 | | |
| | 28 PFG | | | | | | 245 | | |
| | 28 PFG MHG | | | | | | 38 | | |
| | 28 MHG | | | | | | 77 | | |
| | 28AFG MHG | | | | | | 29 | | |
| | 28 AFG PFG MHG | | | | | | 116 | 778 | |
| | 48 AFG | | | | | | 54 | | |
| | 48 AFG PFG | | | | | | 35 | | |
| | 48 PFG | | | | | | 95 | | |
| | 48 PFG MHG | | | | | | 11 | | |
| | 48 MHG | | | | | | 70 | | |
| | 48 AFG MHG | | | | | | 17 | | |
| | 48 AFG PFG MHG | | | | | | 61 | 343 | |
| | 28 48 AFG | | | | | | 80 | | |
| | 28 48 AFG PFG | | | | | | 110 | | |
| | 28 48 PFG | | | | | | 165 | | |
| | 28 48 PFG MHG | | | | | | 25 | | |
| | 28 48 MHG | | | | | | 75 | | |
| | 28 48 AFG MHG | | | | | | 33 | | |
| | 28 48 AFG PFG MHG | | | | | | 92 | 580 | 1,701 |
| **Only overlap H3K27ac peaks** | AFG | | | | | | 684 | | |
| | AFG PFG | | | | | | 294 | | |
| | PFG | | | | | | 913 | | |
| | PFG MHG | | | | | | 97 | | |
| | MHG | | | | | | 484 | | |
| | AFG MHG | | | | | | 173 | | |
| | AFG PFG MHG | | | | | | 654 | | 3,299 |
| **Don't overlap sox17E>sox17M DARs nor H3K27ac peaks** | | | | | | | 107,354 | | 107,354 |
| | | 5,206 | 4,265 | 2,705 | 3,120 | 2,052 | | | **118,088** |

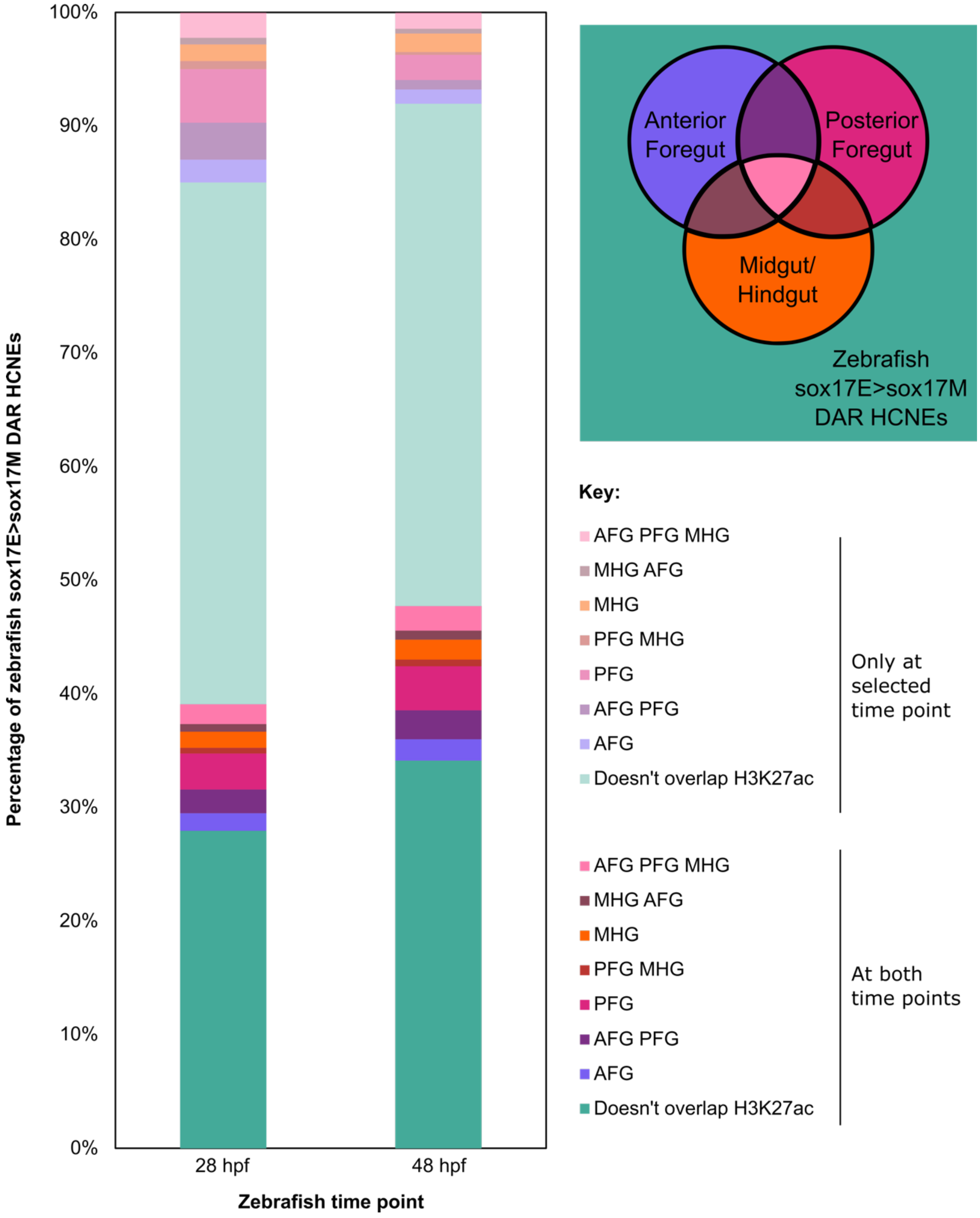


***Supplemental Figure 7. Overlap of HCNEs with sox17E>sox17M DARs and H3K27ac***
*Bar chart to show what percentage of HCNEs that overlap zebrafish sox17E>sox17M DARs at 28 or 48 hpf also overlap human H3K27ac in anterior-posterior patterned endodermal cells and/or zebrafish DARs at the other time point. Venn diagram on the right shows colours used for bar chart with reference to which human endodermal cell population the H3K27ac signal comes from that overlaps the zebrafish sox17E>sox17M DAR HCNEs. Zebrafish sox17E>sox17M DAR HCNEs that do not overlap human endodermal cell population H3K27ac signal are shown in teal.*

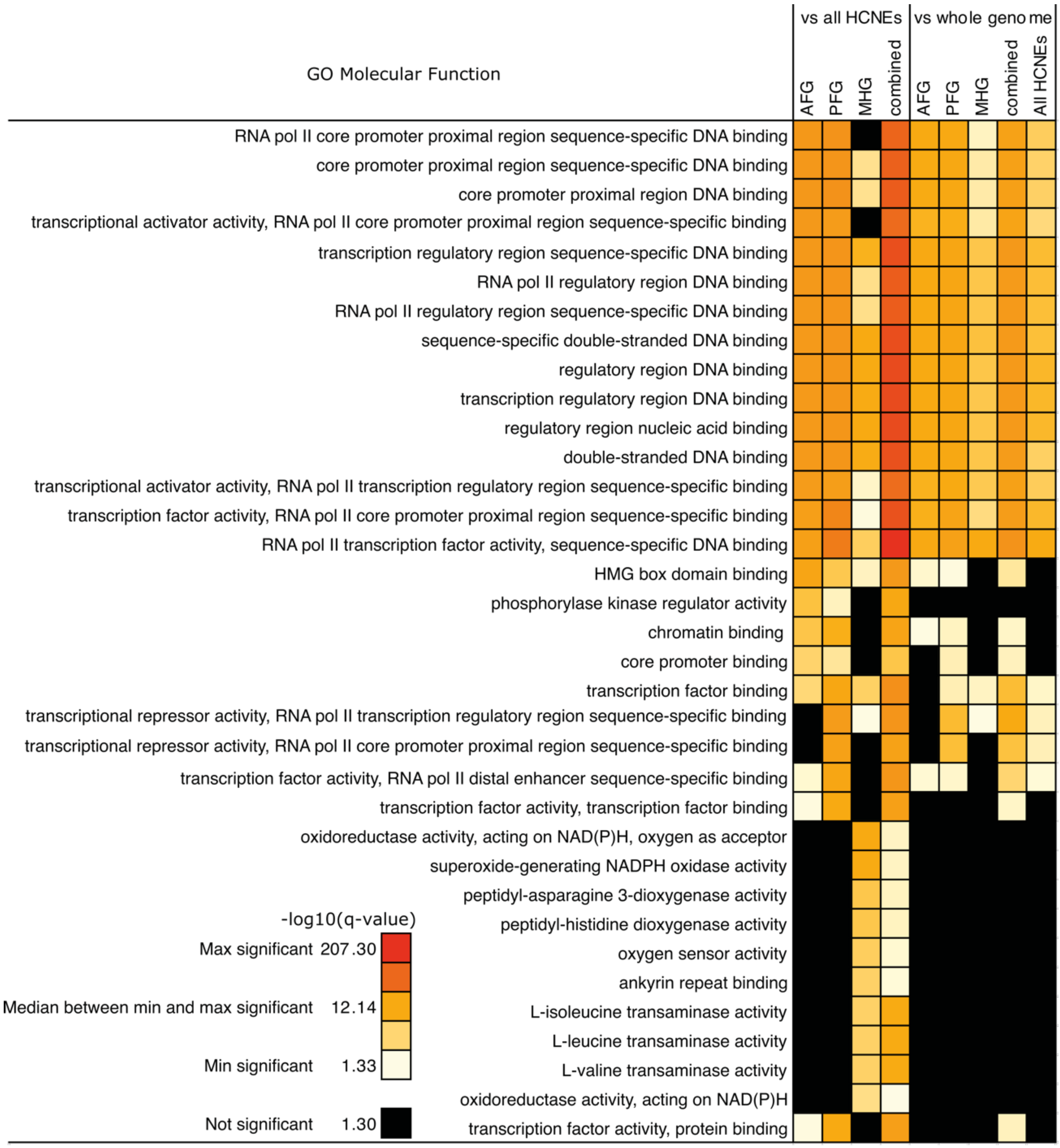


***Supplemental Figure 8. DNA binding terms are found in top GO molecular function terms enriched in genes near endodermal HCNEs***

*HCNEs overlapping sox17E>sox17M zebrafish DARs and H3K27ac peaks in anterior-posterior patterned endoderm cell populations derived from hESCs was used as GREAT input. Terms ranked based on highest -log10(HyperFdrQ). Top 20 terms enriched vs all HCNEs for AFG only, PFG only, MHG only and AFG/PFG/MHG combined were selected and are displayed in that order, with replicates only shown once. Heatmap based on the -log10(HyperFdrQ). Not significant values are shown in black and have a HyperFdrQ<=0.05 and -log10(HyperFdrQ) <= 1.30. Significant values are coloured on a scale from minimum significant -log10(HyperFdrQ) in cream, to maximum significant -log10(HyperFdrQ) in red. The median -log10(HyperFdrQ) value between minimum and maximum significant -log10(HyperFdrQ) is orange.*

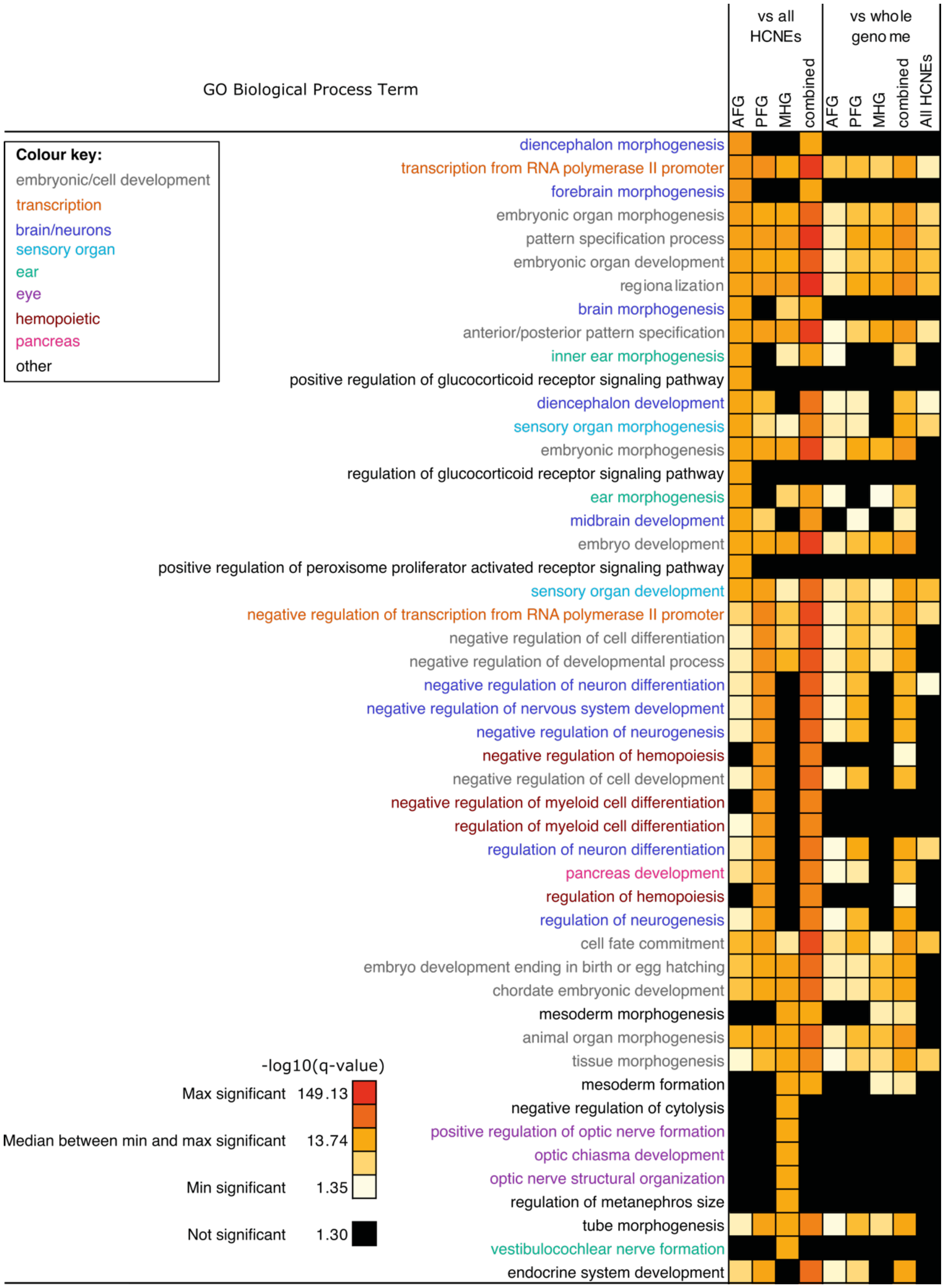


***Supplemental Figure 9. There is enrichment for embryonic development and transcription regulation in the top GO biological process terms enriched in genes near endodermal HCNEs***
*Processed as in (Supplemental Figure 8). Related terms have been coloured as shown in the colour key.*

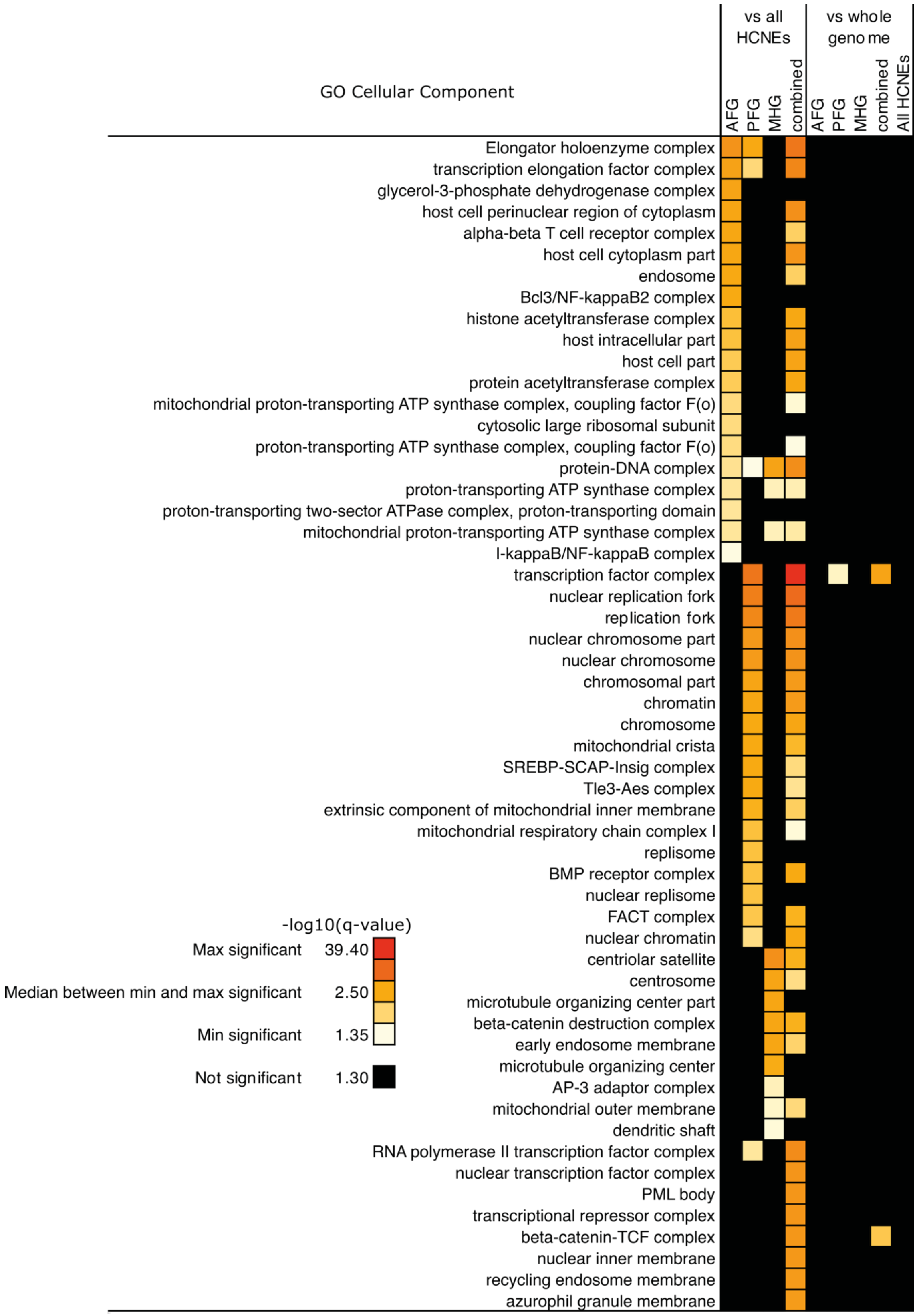


***Supplemental Figure 10. There is enrichment for transcription terms in the top GO cellular component terms enriched in genes near endodermal HCNEs.*** *Processed as in (Supplemental Figure 8).*

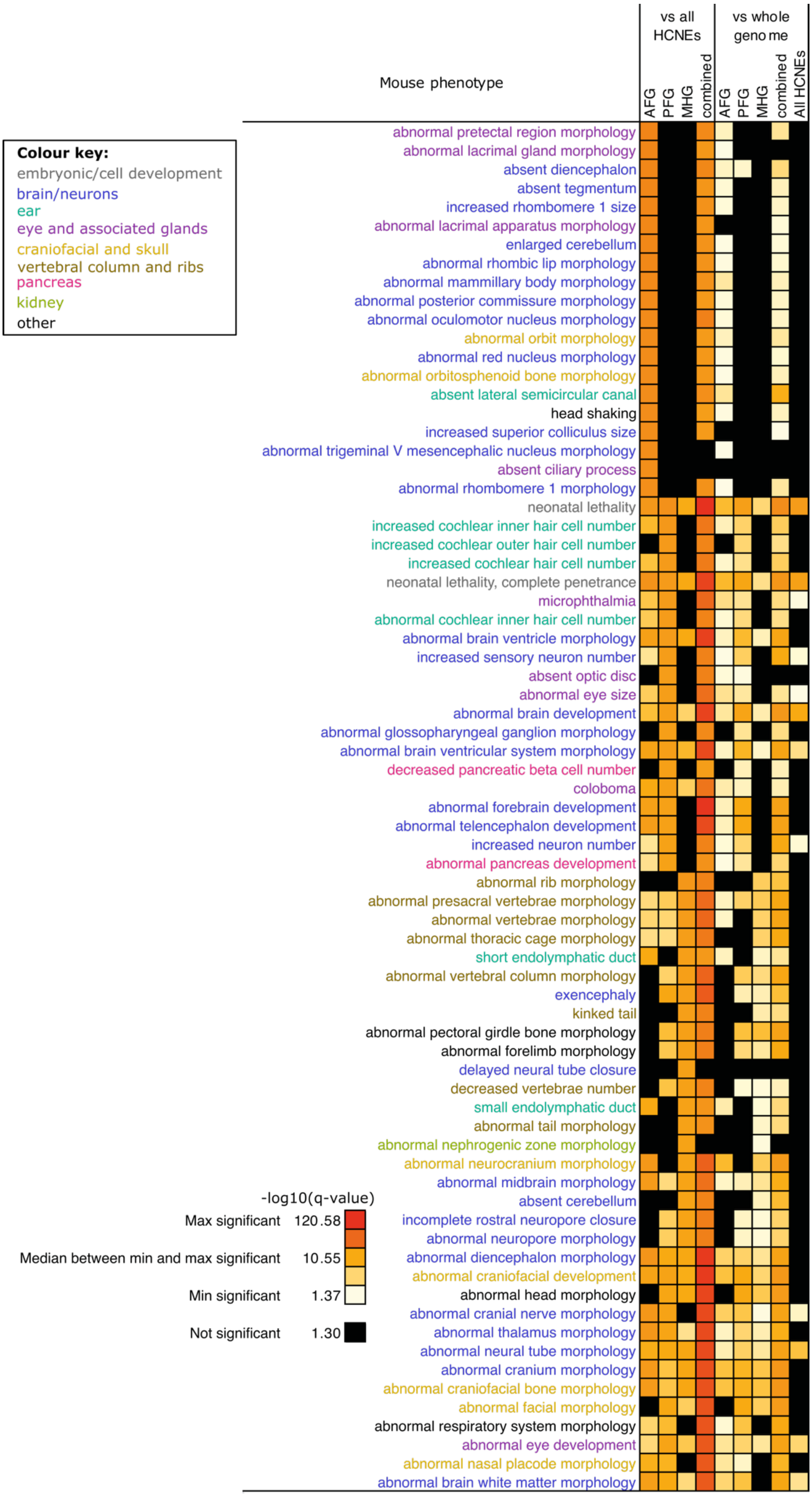


***Supplemental Figure 11. A range of mouse phenotypes are enriched in endodermal HCNE associated genes.*** *Processed as in (Supplemental Figure 8). Related terms have been coloured as shown in the colour key.*

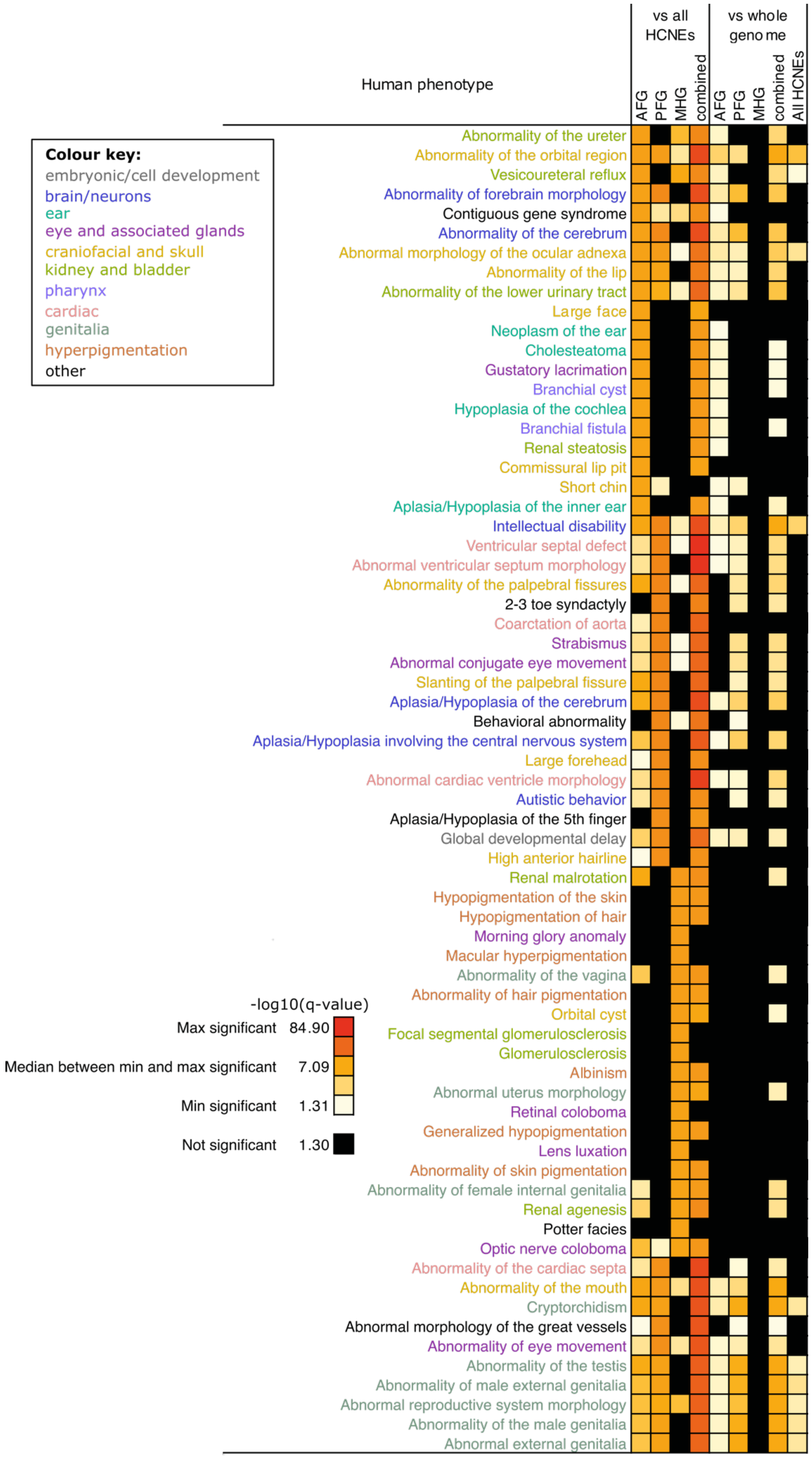


***Supplemental Figure 12. There is enrichment for a variety of phenotypes in the top human phenotype terms enriched in genes near endodermal HCNEs.*** *Processed as in Supplemental Figure 8. Related terms have been coloured as shown in the colour key.*

***Supplemental Table 14. Coordinates of putative enhancers and HCNEs of hnf1ba studied***

| Enhancer name | Chromosome coordinates in genome versions | | | |
|---|---|---|---|---|
| | **danRer7** | **danRer10** | **danRer11** | **hg19** |
| I5Enh +9Kb | chr15:15021553-15022991 | chr15:16066516-16067954 | chr15:16002538-16003976 | chr17:36061008-36070674 |
| I5zHCNE | chr15:15022306-15022646 | chr15:16067269-16067609 | chr15:16003291-16003631 | chr17:36070162-36070510 |
| I5hHCNE | chr15:15022306-15022725 | chr15:16067269-16067688 | chr15:16003291-16003710 | chr17:36070108-36070589 |
| I4 Enh +8Kb | chr15:15023504-15024371 | chr15:16068467-16069334 | chr15:16004489-16005356 | Does not map |
| I4 Enh +6Kb | chr15:15025324-15026283 | chr15:16070287-16071246 | chr15:16006309-16007268 | chr17:36087496-36091612 |
| I4 Enh +6-8Kb | chr15:15023504-15026283 | chr15:16068467-16071246 | chr15:16004489-16007268 | chr17:36087174-36091612 |
| I4zHCNE | chr15:15024917-15025560 | chr15:16069880-16070523 | chr15:16005902-16006545 | chr17:36087165-36087758 |
| I4hHCNE | chr15:15024917-15025560 | chr15:16069880-16070523 | chr15:16005902-16006545 | chr17: 36086990-36087861 |
| Enh -3Kb | chr15:15034801-15035669 | chr15:16079764-16080632 | chr15:16015786-16016654 | Does not map |

***Supplemental Table 15. Table showing where enhancers drove reporter expression in developing zebrafish at 48 hpf***

*For each construct injected the number of embryos (first column) and percentage of embryos screened (second column) that show the observed expression are shown. Intensity of magenta represents percentage of embryos screened showing expression. N refers to biological replicates of experiment, n refers to number of embryos screened. Not scorable embryos are due to there being no expression observed and/or masking of signal due to YSL autofluorescence.*

| Expression observed | **Enhancer construct injected** | | | | | | | | | | | | | | | |
|---|---|---|---|---|---|---|---|---|---|---|---|---|---|---|---|---|
| | **i5** | | | | **i4** | | | | | | | | | | | |
| | **zHCNE** | | **hHCNE** | | **Enh +6-8Kb** | | **Enh +8Kb** | | **Enh +6Kb** | | **zHCNE** | | **hHCNE** | | **Enh -3Kb** | |
| Hindbrain | 2 | 28.6 | 8 | 88.9 | 4 | 28.6 | 0 | 0.0 | 3 | 42.9 | 28 | 62.2 | 14 | 87.5 | 0 | 0.0 |
| Forebrain | 0 | 0.0 | 4 | 44.4 | 0 | 0.0 | 1 | 14.3 | 0 | 0.0 | 1 | 2.2 | 0 | 0.0 | 0 | 0.0 |
| Mouth/jaw | 3 | 42.9 | 0 | 0.0 | 0 | 0.0 | 0 | 0.0 | 0 | 0.0 | 0 | 0.0 | 0 | 0.0 | 4 | 26.7 |
| Neural tube | 0 | 0.0 | 0 | 0.0 | 7 | 50.0 | 6 | 85.7 | 0 | 0.0 | 3 | 6.7 | 1 | 6.3 | 0 | 0.0 |
| Floor plate | 0 | 0.0 | 0 | 0.0 | 5 | 35.7 | 0 | 0.0 | 0 | 0.0 | 1 | 2.2 | 0 | 0.0 | 0 | 0.0 |
| Pericardium and/or heart | 4 | 57.1 | 3 | 33.3 | 0 | 0.0 | 0 | 0.0 | 0 | 0.0 | 0 | 0.0 | 0 | 0.0 | 0 | 0.0 |
| Blood | 0 | 0.0 | 4 | 44.4 | 0 | 0.0 | 1 | 14.3 | 0 | 0.0 | 0 | 0.0 | 0 | 0.0 | 0 | 0.0 |
| Tail muscle cells | 0 | 0.0 | 2 | 22.2 | 0 | 0.0 | 4 | 57.1 | 0 | 0.0 | 3 | 6.7 | 1 | 6.3 | 0 | 0.0 |
| Not scorable | 0 | 0.0 | 1 | 11.1 | 0 | 0.0 | 0 | 0.0 | 3 | 42.9 | 17 | 37.8 | 1 | 6.3 | 0 | 0.0 |
| Number of embryos screened (n) | 7 | 100.0 | 9 | 100.0 | 14 | 100.0 | 7 | 100.0 | 7 | 100.0 | 45 | 100.0 | 16 | 100.0 | 15 | 100.0 |

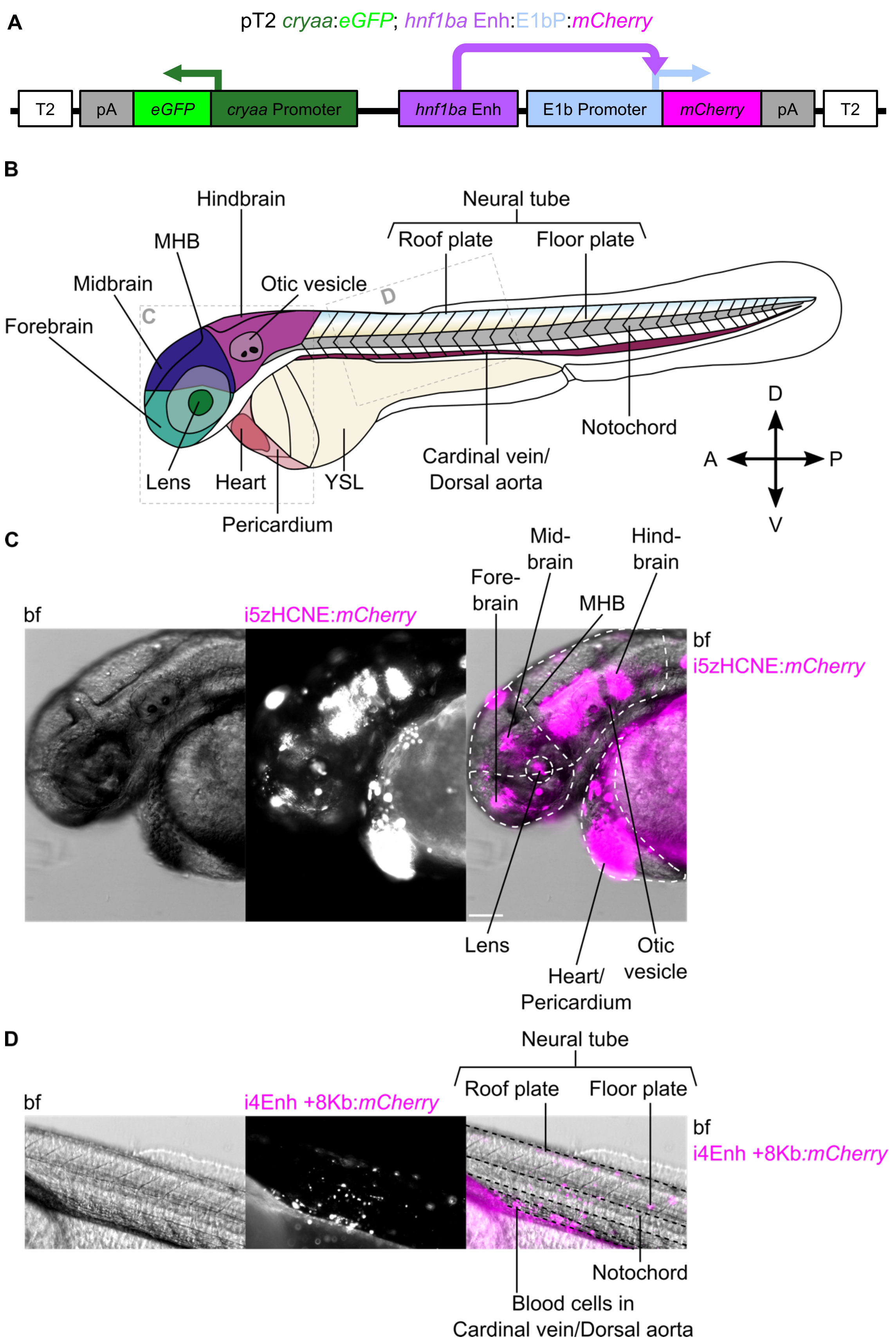


***Supplemental Figure 13. Putative hnf1ba enhancers show reporter expression in multiple cell types including in the brain and neural tube***

***A.*** *Schematic of reporter construct injected alongside Tol2 mRNA for reporter assay. Green arrow indicates cryaa driving eGFP expression. Purple arrow indicates putative enhancer enhancing mCherry expression. Blue arrow indicates E1b promoter driving mCherry expression.* ***B.*** *Schematic of expression domains seen in zebrafish at ~48 hpf. Gray boxes with letters denote the approximate location of the images shown in C and D. For the anatomical direction: D = dorsal, P = posterior, V= ventral, A = anterior. MHB = midbrain hindbrain boundary. YSL = yolk syncytial layer.* ***C-D.*** *Images from 48 hpf embryos injected with reporter constructs to demonstrate some of the expression patterns seen. Left is bf, middle is reporter with basal promoter E1b driving mCherry expression, and right is overlay of bf and mCherry (magenta). Lateral views. Scale bar = 100mm. Contrast settings are the same for both C and D. Dashed lines show approximate boundaries for anatomical structures labelled.* ***C.*** *i5zHCNE driving expression in forebrain, midbrain, hindbrain, and heart/pericardium.* ***D.*** *i4Enh +8Kb driving expression in the roof plate and floor plate of the neural tube, as well as the blood cells in the cardinal vein/dorsal aorta.*

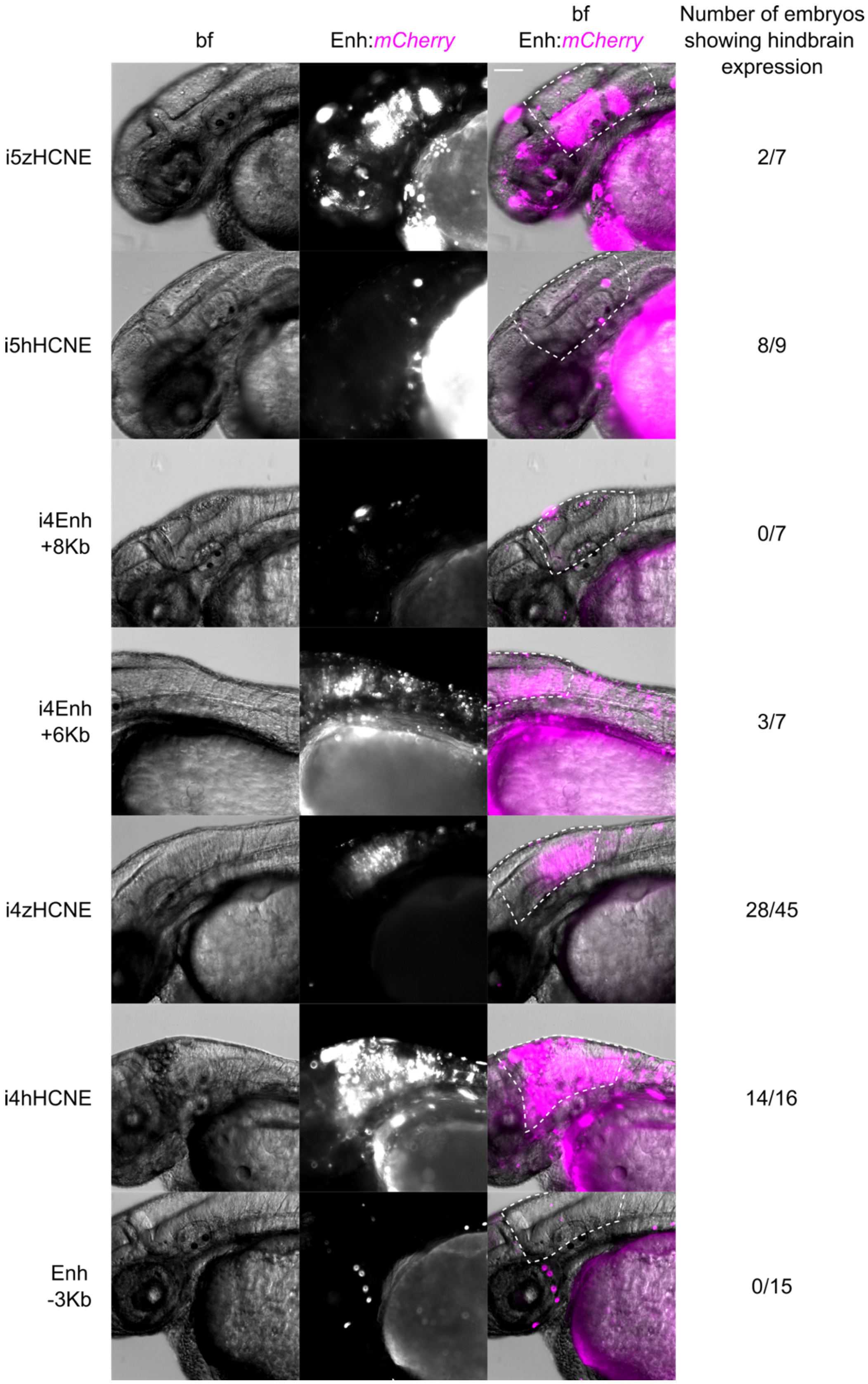


***Supplemental Figure 14. Consistent expression in the hindbrain driven by hnf1ba putative enhancers.*** *Images of heads of 48 hpf embryos injected with reporter construct. Lateral views. All are dorsal up, and anterior on the left (i4zHCNE has been flipped horizontally for ease of comparison). Left is bf, middle is enhancer with basal promoter E1b driving mCherry expression, and right is overlay of bf and mCherry (magenta). White dashed line marks hindbrain. Scale bar = 100mm. Contrast settings are the same.*

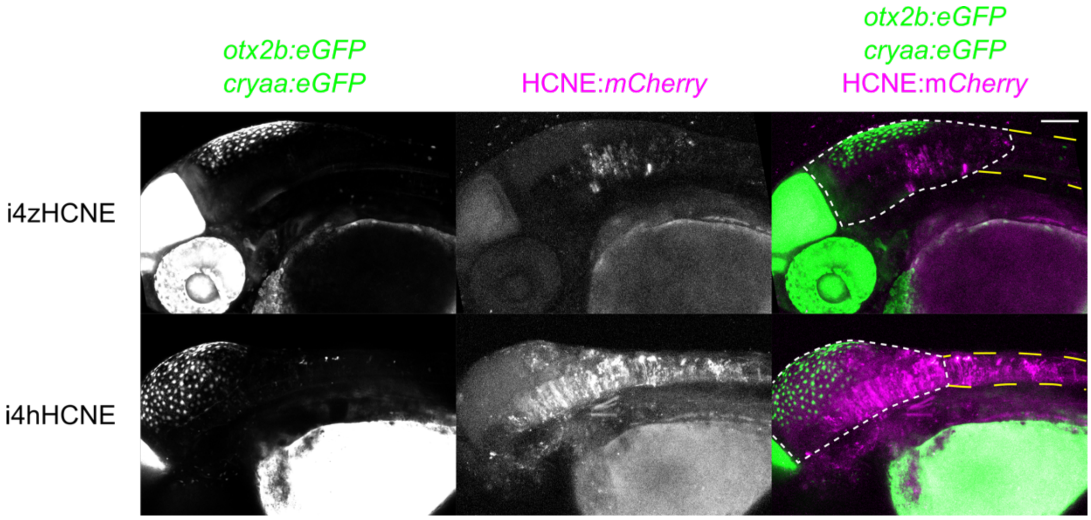


***Supplemental Figure 15. Hnf1ba HCNEs from human and zebrafish both drive expression in the hindbrain.*** *48 hpf Tg(otx2b:EGFP) embryos injected with reporters i4zHCNE (top) and i4hHCNE (bottom). Left is otx2b:EGFP and cryaa:EGFP, middle is reporter with basal promoter E1b driving mCherry expression, and right is overlay of eGFP (green) and mCherry (magenta). 27 confocal z-slices were merged by z-projection based on maximum intensity. Embryos are orientated laterally. Dorsal is up, and anterior is on the left. Short white dashed line marks hindbrain, long yellow dashed line marks neural tube. Scale bar = 100mm. Embryos are representative of embryos that have the reporter construct incorporated (have cryaa:EGFP expression in lenses) and show expression of the HCNE in the hindbrain.*

***Supplemental Table 16. Primers used in this study***

***Primers to amplify putative hnf1ba enhancers from genomic DNA***

| Primer name | Sequence (5' to 3') Forward | Reverse |
|---|---|---|
| I5Enh +9Kb | ggggacaactttgtatagaaaagttg ACCAGCCAGTCAGTGTTACA | ggggactgctttttgtacaaacttg ACGACGTCAAGAGGAGAATGT |
| I5zHCNE | ggggacaactttgtatagaaaagttg TAGCAGCAATGCCATGTCAAC | ggggactgctttttgtacaaacttg AGCTACATGAAGATCTCGCC |
| I5hHCNE | ggggacaactttgtatagaaaagttg CAGCCAGTCGGTTTTACAGC | ggggactgctttttgtacaaacttg GAAGAGGGGGCTTGTGTCAA |
| I4 Enh +8Kb | ggggacaactttgtatagaaaagttg TCGCGTTCTCAGAGGTTGTA | ggggactgctttttgtacaaacttg GCTGAGTTTTGTTGTGTAGCG |
| I4 Enh +6Kb | ggggacaactttgtatagaaaagttg AGCAAAGAGGAGAGCAGTGA | ggggactgctttttgtacaaacttg GCAAACAGACAAGCCACTCA |
| I4 Enh +6-8Kb | ggggacaactttgtatagaaaagttg AGCAAAGAGGAGAGCAGTGA | ggggactgctttttgtacaaacttg GCTGAGTTTTGTTGTGTAGCG |
| I4zHCNE | ggggacaactttgtatagaaaagttg GCACTGCAATGAGTGGCTTG | ggggactgctttttgtacaaacttg GACCTTCTAATGGCGGCGAA |
| I4hHCNE | ggggacaactttgtatagaaaagttg CCAAACATCAGCACCTGAGC | ggggactgctttttgtacaaacttg CTGCCGACTAGAGCAAAGGG |
| Enh -3Kb | ggggacaactttgtatagaaaagttg GTGCACGCAACTCTAAGGTT | ggggactgctttttgtacaaacttg TGGGGTGTACTTAATTATGCTGA |

***Primers to switch eGFP for mCherry in reporter constructs***

| Primer name | Sequence (5' to 3') Forward | Reverse | Use |
|---|---|---|---|
| eGFP | gacattaccgATGGTGAGCAAGGGCGAG | ggccgctttaCTTGTACAGCTCGTCCATGC | Amplify *eGFP* region from pENTR bas:EGFP |
| mCherry | GTGAGCAAGGGCGAGGATGACAACATGG | tgagtccggaTTACTTGTACAGCTCGTCC | Amplify *mCherry* region from pDEST tol2 pA *cryaa: mCherry* |
| pENTR baseGFP | gtacaagtaaTCCGGACTCAGATCTCGAG | tcatcctcgcCCTTGCTCACCATGGTGG | Amplify pENTR bas:EGFP backbone without *eGFP* |
| pDESTtol2pA cryaamCherry | gctgtacaagTAAAGCGGCCGCGACTCTAG | tgctcaccatCGGTAATGTCAGACCTGGTAAC | Amplify pDEST tol2 pA *cryaa:mCherry* backbone without *mCherry* |

***Primers to amplify E1b promoter for HiFi insertion into pENTR plasmid***

| Primer name | Sequence (5' to 3') Forward | Reverse |
|---|---|---|
| E1b promoter HiFi | acaaaaaagcaggctcgcta CTCGACTCTAGAGGGTATATAATG | cttgctcaccatggtggcga TTTGCCAAAATGATGAGAC |
